# Epigenetic feedback on noisy expression boosts evolvability

**DOI:** 10.1101/2022.06.29.498068

**Authors:** Werner Karl-Gustav Daalman, Liedewij Laan

## Abstract

Adapting organisms often face fitness valleys, i.e. barriers imposed by ubiquitous genetic interactions, while optimizing functions. Elucidating mechanisms that facilitate fitness valley traversals is integral to understanding evolution. Therefore, we investigated how protein expression noise, mechanistically decomposed into instant variation and epigenetic inheritance of optimal protein dosage (‘transgenerational feedback’), shapes the fitness landscape. For this purpose, we combined a minimal model for expression noise with diverse data of *Saccharomyces cerevisiae* from literature on e.g. expression and fitness to representatively simulate mutational fitness effects. For our proxy of point mutations, which are very often near-neutral, instant dosage variation by expression noise typically incurs a 8.7% fitness loss (17% in essential genes) for non-neutral point mutations. However, dosage feedback mitigates most of this deleterious effect, and additionally extends the time until extinction when essential gene products are underexpressed. Taken together, we consider dosage feedback as a relevant example of Waddington’s canalization: a mechanism which temporarily drives phenotypes towards the optimum upon a genetic mismatch, thereby promoting fitness valley traversal and evolvability.

**Author summary:** Gene products frequently interact to generate unexpected phenotypes. This universal phenomenon is known as epistasis, and complicates step-wise evolution to an optimum. Attempts to understand and/or predict how the optimum is found are further compromised by the countless combinations of mutations that are considered by nature, and necessitate the formulation of general rules on how the obstacles that epistasis presents are bridged. To make such a rule as insightful as possible, we reduced cell division to a generation-based model focusing on one protein at a time for reproductive success. Importantly, protein production between divisions is stochastic and we show how the resulting expression noise affects epistasis. After validating the model on experimental fitness landscapes, we combine high-throughput data of budding yeast from multiple sources to make our model predictions on mutational effects on fitness as representative as possible. We find different effects per mutation type: gene duplications have little effect, as genes in our simulated pool are rarely toxic, loss-of-function mutations decrease mutational gains as adaptation progresses, and point mutations permit expression noise to unlock its roles in adaptation. For non-neutral point mutations, noise imposes a sizeable fitness penalty or even induces extinction, which is alleviated by an epigenetic, transgenerational feedback on protein dosage which is never deleterious. Particularly for essential genes, we predict that this effect reduces the obstacles of epistasis and hence significantly increases evolvability, adding to the general rules of evolution.

## Introduction

The ability to predict evolution has a plethora of societal applications. For example, tracking and forecasting viral evolution benefits vaccination strategies (Du, King, Woods, & Pascual, 2017; Neher & Bedford, 2015), efforts in rational design and directed evolution of microbial communities will improve chemical production and many aspects of daily life (Sanchez et al., 2020; Zomorrodi & Segrè, 2016), and research on evolutionary models provides better grip on diseases such as cancer (Diaz-Uriarte & Vasallo, 2019). However, several confounding phenomena complicate our understanding of evolution. One of those is expression noise, i.e. variation in dosage of gene products across an isogenic population. Another example is epistasis (e.g., (Bank, Matuszewski, Hietpas, & Jensen, 2016; Miton & Tokuriki, 2016; Sailer & Harms, 2017)), i.e. the observation that mutational effects can depend on the particular genetic background in terms of magnitude and sign, which causes the adaptive fitness landscape to have non-trivial shapes. Here we ask how expression noise and epistasis mechanistically intertwine, to ultimately improve our understanding and accurate prediction of evolution.

Previous research has primarily focused on the consequences of noise given a certain fitness landscape, with ambiguous conclusions. On the one hand, noise can drive a population away from the optimal equilibrium, while on the other hand, noise can also function as a bet-hedging strategy (Philippi & Seger, 1989). This ambiguity also holds for the particular case of expression noise. Theoretically, expression noise can influence environmental robustness (Mineta, Matsumoto, Osada, & Araki, 2015), but it can also decrease the effective population size and increase drift (Wang & Zhang, 2011). By the same token, there is empirical evidence that expression noise has both been selected for as well as selected against in the evolution of *Saccharomyces cerevisiae* (Fraser, Hirsh, Giaever, Kumm, & Eisen, 2004; Z. Zhang, Qian, & Zhang, 2009).

Equally interesting is how noise shapes the fitness landscape itself. While theoretically important (Coomer, Ham, & Stumpf, 2022), the empirical relevance of this noise interaction has been much less intensively studied thus far. Epistasis is a natural property of fitness landscapes for noise to interact with, as it is universally found across many organisms (Sanjuán & Elena, 2006). For example, changes in expression are expected to be a common source of (sign) epistasis (Li, Lalić, Baeza-Centurion, Dhar, & Lehner, 2019). This epistasis (see conceptually in Figure 1A) is particularly important for essential genes. As essentiality is increasingly found to be context-dependent (Larrimore & Rancati, 2019), epistasis and its link to noise are highly relevant for adaptation. Another example of ubiquitous negative epistasis is diminishing returns, i.e. mutations becoming less beneficial as fitness increases, which has been shown in model systems such as *Methylobacterium extorquens* (Chou, Chiu, Delaney, Segrè, & Marx, 2011), *Escherichia coli* (Khan, Dinh, Schneider, Lenski, & Cooper, 2011), *S. cerevisiae* (Kvitek & Sherlock, 2011) and a multicellular fungus (Schoustra, Hwang, Krug, & de Visser, 2016). The generality of diminishing returns epistasis across unrelated biological modules (Kryazhimskiy, Rice, Jerison, & Desai, 2014) naively suggests the existence of generic causes. As variation in protein dosage is known to couple otherwise unrelated modules (Kleijn, Krah, & Hermsen, 2018), a possible hypothesis is that expression noise generically affects epistasis and thereby the shape of the fitness landscape.

**Figure 1.**
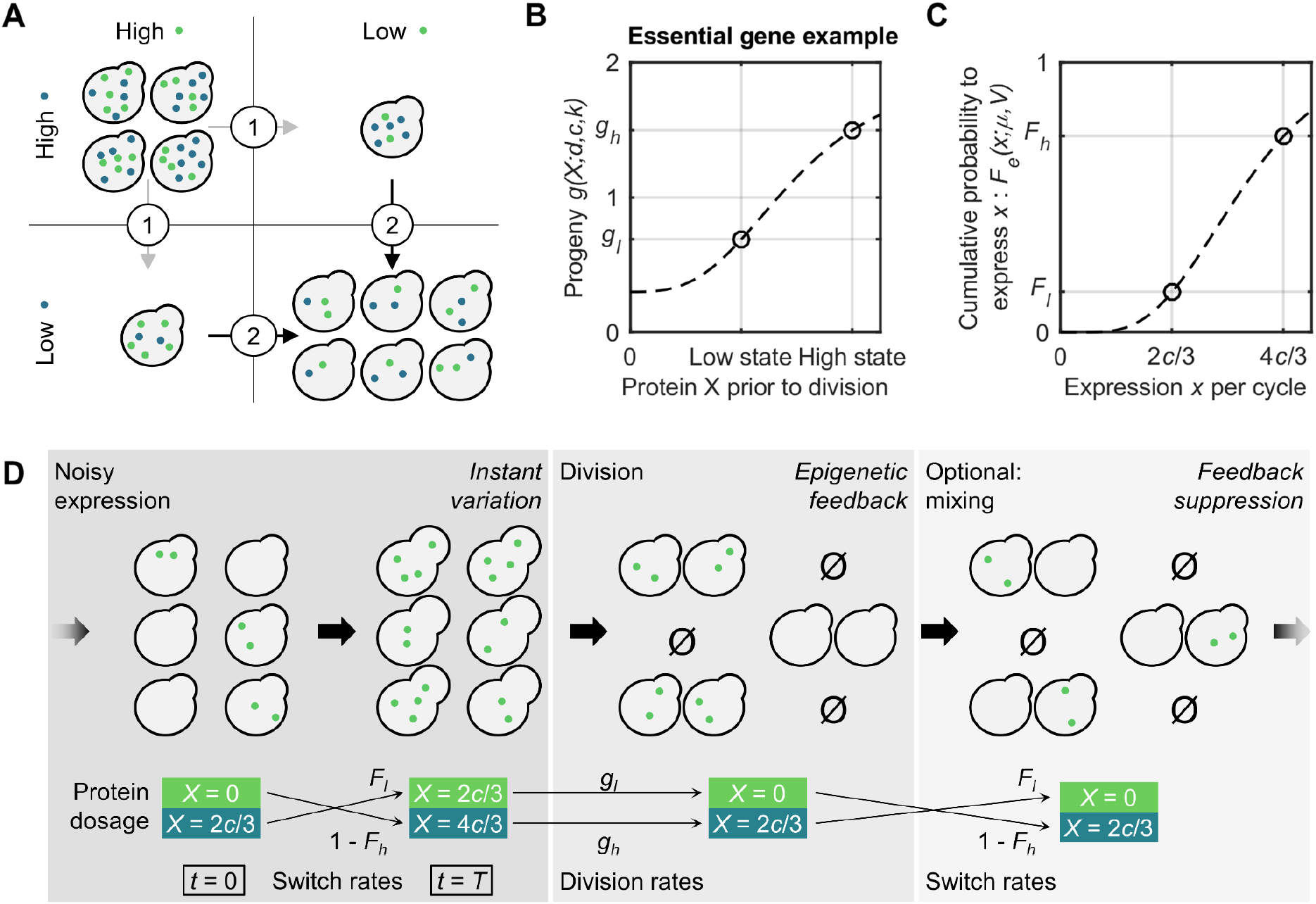
Overview of the generation-based MEN-population model for adaptive epistasis under influence of expression noise. (A) Conceptual view of (sign) epistasis between two genes, whose gene products can be in a low or high dosage. Starting at high dosages, the first adaptive step always reduces fitness, but must precede the second step to yield an overall superior fitness. The number of cells in this cartoon depicts the fitness of that state. (B) Model description of progeny *g* from an individual cell as function of a single protein dosage *X* following a Hill curve, in this case illustrated for an essential gene product. Two states symmetric to the tipping point concentration *c* in the curve can be identified: low or high dosage. The progeny in the worst state is defined as *g_l_*, in the best state as *g_h_*. Parameters *d* and *k* determine the depth and steepness of the Hill curve. (C) When protein is expressed in one burst per cell cycle (in amount *x*), cells can switch states stochastically each cycle with probabilities following the cumulative distribution function *F_e_* to switch dosage states. *F_l_* and *1-F_h_* are the respective chances to switch from the high to low state or vice versa. Probabilities depend on mean expression level *μ* and expression coefficient of variation (noise level) *V*. (D) Graphical overview of the minimal epistasis-noise (MEN-) model concerning a population of cells that once per cycle time *T* can stochastically switch between two protein dosage states (high/low, represented by dots) before division. The right-most patch denotes an optional addition to the MEN-model, where inheritance of dosage is suppressed by resetting the population dosage distribution.

In this paper, we will study how the interaction between noise and fitness landscapes feeds back on epistasis, differentiating between genetic and epigenetic contributions of noise. For this purpose, we construct a minimal cell model to understand how noise shapes fitness landscapes and mutational effects therein, decomposed by mutational type and noise mechanism. This minimal model focusses on a single gene product stochastically switching between two dosage states, which determine the progeny per cell. First, we explore theoretically how the evolutionary roles of noise relate to two distinct noise mechanisms, namely generating instantaneous variation in protein dosage and an epigenetic feedback on dosage. Then, we integrate literature data of *S. cerevisiae* on for example empirical fitness landscapes and the distribution of mutational effects, to generate a representative estimate of the pattern of epistasis for e.g. promoter mutations and duplications. We find that an simple epigenetic feedback biases dosage with every generation to cause expression noise to significantly mitigate the fitness penalties associated with instant variation, particularly for essential genes. Furthermore, the feedback also promotes longer survival before extinction when gene products are on average at a lethal dosage. We conjecture that this noise-based epigenetic feedback is important for the evolvability of many essential genes across organisms, suggesting a general rule on how expression noise acts as a fitness landscaper. Consequently, feedback emerges as a canalization mechanism in the context of evolutionary theories.

## Model methods

### Construction of a minimal model for epistasis and noise

To elucidate the mechanistic coupling of noise to fitness landscapes, we make a minimal model for epistasis subject to noise (MEN-model), depicted in Figure 1D. The minimality of this model has the benefit of tractability and also accommodates the seemingly generic nature behind epistasis-noise coupling by disregarding many biological details. The latter justifies coarse-graining the underlying protein interactions and reducing the cellular environment to one protein under consideration at a time.

We model a population of cells containing a protein X, with fitness *ω* defined as the reciprocal of the population doubling time (see also Model implementation details). In short, we disregard degradation and dilution, and the constant cell size permits interchangeable use of protein number and concentration. The cell cycle is simplified to two stages: first, X is produced stochastically, proportional to the cell cycle duration *T*. After this time, each cell is replaced by two cells which inherit X evenly, of which the progeny number *g* (between 0 and 2) survive. We assume the progeny scales with the available concentration [X], which is parametrized as a Hill curve (for example as in Figure 1B), where *d* modulates the difference between the best and worst progeny state:

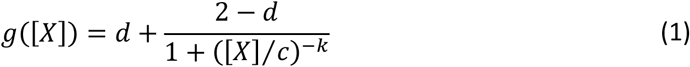

The Hill equation loosely permits the generic interpretation of X being involved in cooperative binding (with some caution (Prinz, 2010)) or switching between activation states (e.g., by phosphorylation). This defines *c* as a tipping point concentration and |*k*| as an effective cooperativity coefficient, resulting from coarse-graining the full chemical pathway involving X.

After division, X is binned into two dosages, either low or high with respect to *c*, and therefore we only evaluate the progeny function *g* at 2*c*/3 and 4*c*/3, with values defined as *g_l_* and *g_h_* respectively. Consequently, the stochastic production of X forms a two-state discrete time stationary Markov chain, where every time step is a generation. Coefficients in the transition matrix *M* denote switching probabilities following from the cumulative distribution function (cdf) *F*_e_(x; *μ, V*) governing expression, with x as the added protein and the parameters *μ* as mean expression and *V* as noise level. We only require the cdf at *x* = 2*c*/3 and 4*c*/3, whose values we define as *F_h_* and *F_l_* respectively (see Figure 1C). This yields state vector *f*, whose entries represent the number of cells in the high state (*f_h_*) or low state (*f_l_*) respectively:

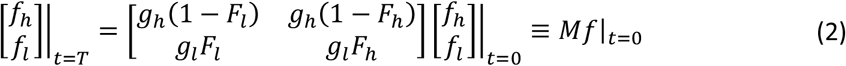

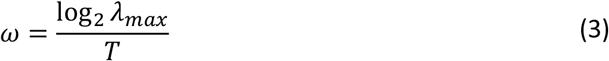

where *λ_max_* is the largest eigenvalue of *M* and *ω* is the fitness at equilibrium, which we can multiply by *T* to obtain the relative fitness ω_Γ_. Conveniently, our fitness is in reasonable accordance with the fit function postulated in (Keren et al., 2016), so we can straightforwardly compare the two fit results (see MEN-model fitness comparison with literature). We note that our model provides a parsimonious and interpretable alternative on the originally postulated fit function, improving on 84% of the landscape data (see Appendix 1-figure 1).

### Decomposition of noise into variation and feedback

We distinguish two noise contributions. Firstly, instant variation generates different dosages within each cycle. Additionally, as some cells have a dosage that allows more progeny, and since dosage is heritable, this results in an epigenetic bias towards the favorable dosage unless protein life-times are very short. We can refer to this as a transgenerational feedback of protein dosage after (Xue & Leibler, 2016), only with respect to a genetic rather than a physical environment.

We implement the *absence* of feedback into the model by resetting the distribution of cells across the two states after every division (see Figure 1D), countering the effect of selection. We redistribute the cells across the two states according to the state proportions when selection is absent, so when *g_h_* = *g_l_* = 2 in equation 2. This feedback has a strictly non-negative effect (see Appendix) on fitness or exponential decay of the population (λ_*max*_ < 1). The relative fitness is given by:

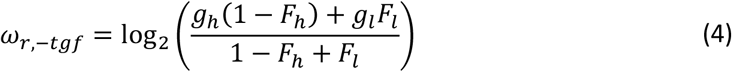

### Generation of distributions of fitness effects (DFEs)

To simulate the effects of mutations on fitness in populations in the MEN-model, we incorporate mutations into the model in two ways. Firstly, we consider expression mutations that affect the mean of the distribution underlying *F*_e_(x; *μ, V*). Mutations that affect fitness otherwise are incorporated as a generic effect on cycle time *T*. Because a change in cell cycle time *T* also affects mean expression, these generic mutations can couple to expression in an unrelated module, akin to (Kleijn et al., 2018). Combining these mutation types, we examine the distribution of fitness effects (DFEs) by expression mutations in different stages of adaptation by varying the cycle time through the generic mutations.

It is possible to link the simulated mutations considered above to many concrete mutations, which induce an effective change in mean expression. Apart from the trivial case of synonymous substitutions, this category includes promoter mutations, duplications, deletions or premature stops, mutations that affect reaction rates involving protein X that lead to an effective change in amount of available X, mutations that change mRNA stability, mRNA lifetime or protein half-life, but also 5’UTR mutations and RBS mutations influence expression (Kosuri et al., 2013; Mutalik et al., 2013). Expression related mutations form a significant part of the adaptive mutations in evolutionary trajectories of *S. cerevisiae* (Kryazhimskiy et al., 2014). It is even plausible that our modelled expression related mutations encompass synonymous mutations that affect fitness (Shen, Song, Li, & Zhang, 2022).

## Results

### Noise theoretically delays extinction, feedback compensates fitness loss

In order to provide more intuition about possible fitness landscapes, we first formulate some expectation based on a theoretical analysis of the MEN-model, before applying the model to realistic mutational scenarios. In Figure 2, we consider the two extreme situations: the essential (lethal when underexpressed) and toxic (lethal when overexpressed) landscapes. We assume very sharp Hill curves (high effective cooperativities, *k*>>1) in both cases, and an initial population size of 1 million which is only relevant when the population is exponentially decaying. Three roles of noise immediately emerge, which we can decompose into the two aspects of noise: instant variation and epigenetic feedback.

**Figure 2.**
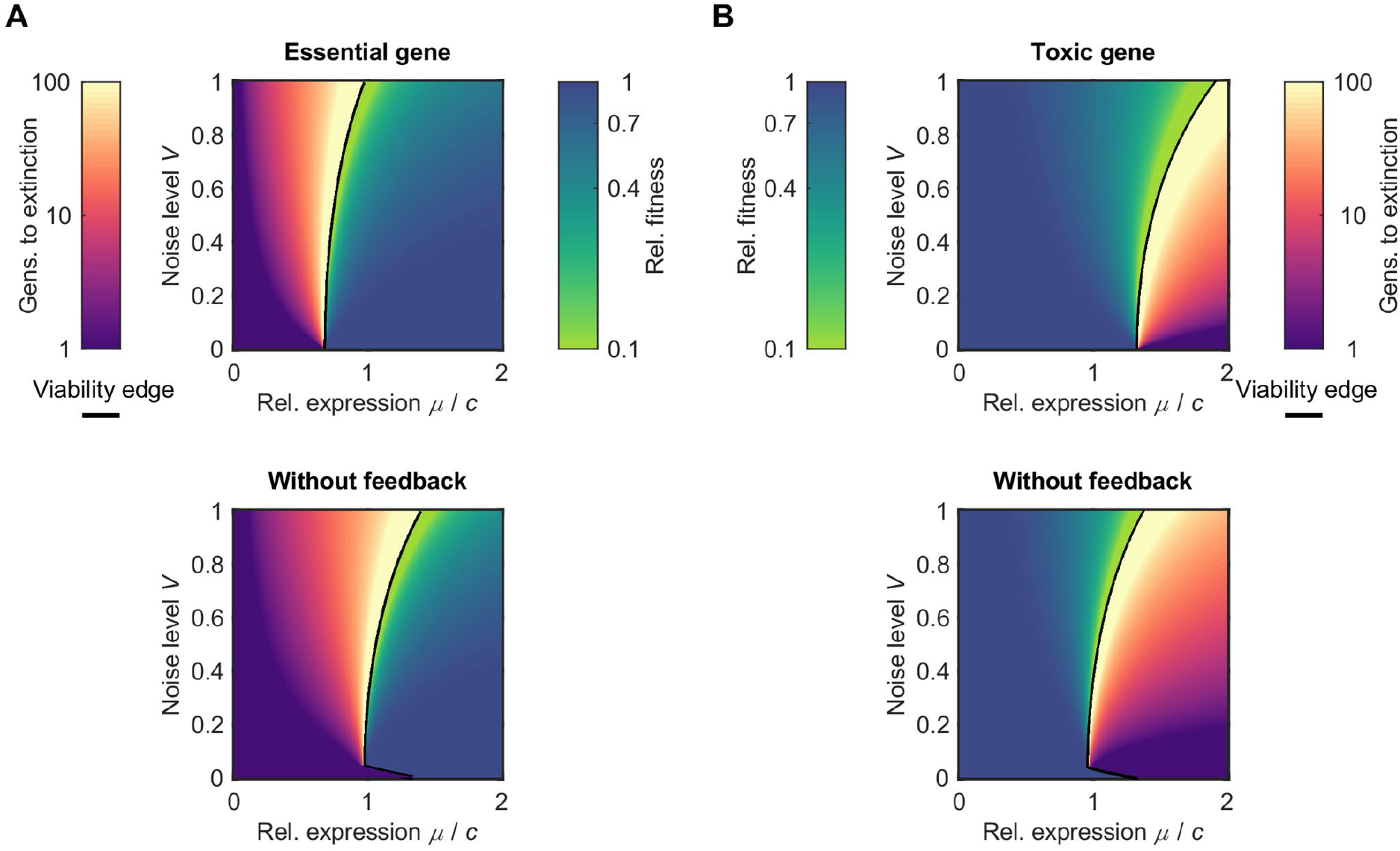
Two fitness landscape types with structurally and temporarily viable regimes, with and without feedback. Landscapes are given as heat maps plots as function of relative expression *μ* (scaled by tipping point concentration *c*) and noise level *V*. Blue to light green colors denote the log of relative fitness *ω_r_*, whereas the yellow to purple colors denote how many generations a population of 10^6^ cells is expected to survive. The two regimes are separated by a black line. All plots assume a gamma cdf *F_e_*, essential and toxic genes have | *k*|=10^5^ and *d*=0. For completeness, noise levels below the feasible range in (Chong et al., 2015), where 99.8% of genes falls in 0.1<V<1, are plotted, with values below the discontinuity in the viability edge approximated. Color maps from (Smith, Walt, & Firing, 2015).

Firstly, noise smoothens the fitness landscape. For expression levels where the fitness can be defined (blue to green colors), increasing noise increases the span of the color gradient. Analogously, the number of generations a population can temporarily survive (red to purple colors) when expression is structurally insufficient or too high, also exhibits a shallower gradient at higher noise levels. We see this smoothing effect with and without feedback, although it is theoretically expected to be slightly more pronounced with feedback unless the landscape is relatively flat (see Figure 2-figure supplement 2).

Secondly, increased noise has a dual effect on fitness. For the essential gene landscape, we see fitness decrease with the addition of noise, in correspondence with the notion that noise drives the population away from equilibrium. By contrast, in the toxic gene landscape, noise can actually improve fitness. However, we will see later that in practice, this beneficial role of noise will be rare. The feedback has no effect on the reversal from a negative to positive effect on fitness depending on landscape shape. The cause for this reversal is instant variation.

Thirdly, we see a widespread role for noise to improve temporary survival. When expression is insufficient to sustain the population, population size will instead shrink exponentially with every generation. The number of generations it takes for the population to go extinct is important for evolution. For example, an environmental perturbation may render a population with a certain genotype inviable, causing a population shrinkage with time/generations. In the case of budding yeast, a large inviable population may also arise from sporulation. As the population decays, it can only evade extinction if it finds a genetic mutation that returns the population to the viable expression regime. We see noise conveniently increases the number of generations until extinction and thereby the time available for a compensatory mutation to occur. The diversifying and buffering potential of noise is not released in one moment, but is replenished continuously. Again, we see this effect with and without feedback.

Despite qualitative similarities between scenarios with and without feedback, an important function surfaces for the transgenerational feedback on protein dosage. Comparing the two plots in the top row of Figure 2 to those in the bottom row, we see that deleting the memory of the system causes several deleterious effects on the population: (i) for expression levels where a fitness can be defined, the fitness is decreased; (ii) the viability edge, the expression threshold for which the population can structurally sustain its survival, also shifts to a disadvantageous direction; (iii) the number of survivable generations beyond the viability edge is decreased. So, although the feedback does not yield qualitatively different behavior for the effect of noise on fitness, there are noticeable quantitative effects on fitness. We reiterate that these quantitative effects of the feedback are never deleterious (see Strict non-negativity of transgenerational feedback effect on fitness). We also note that this feedback requires a minimal amount of noise at any finite population size to sustain dosage memory inside the population in practice, as marked by the discontinuity in e.g. the viability edge. However, given the feasible region in (Chong et al., 2015) of 0.1<*V*<1, we can safely ignore this exception.

### Translation to realistic mutational landscapes

#### Validation of MEN-model fits

After forming theoretical expectations, we validate the MEN-model as part of our translation to realistic mutational scenarios in yeast by evaluating model fits, for estimates of the parameters *k, c, d* and *V* on empirical fitness landscapes (Keren et al., 2016) (see also Materials and Methods section and SI: Evaluation MEN-model fits on empirical fitness landscapes). These landscapes are combined with WT dosage data (Kulak, Pichler, Paron, Nagaraj, & Mann, 2014) and essentiality data (Cherry et al., 2012). Compared to the original fits of (Keren et al., 2016), our model fits are improvements in 84% of the cases (metric R^2^ adjusted for the free parameters (Wherry, 1931)), see Appendix 1-figure 1A.

However, despite the quality of the fits, the interpretation of parameters should be approached with care. The Hill curve for the progeny function suggests the interpretation of *k* as a Hill coefficient, which in turn can represent allostery (e.g., (Prinz, 2010)). Yet, if we define a feasible allosteric range of | *k* | ≤ 5 for simple reactions, that means that many of the inferred *k*’s fall outside this reasonable regime (Appendix 1-figure 1C). This suggests that a generic reaction form involving protein X assumed is probably too simplistic for most gene products considered. The observed ultrasensitivity suggests more subtle underlying processes, such as feedback loops or multiple phosphorylation steps to activate one protein (Ferrell & Ha, 2014a, 2014b; Ferrell, Jr, & Ha, 2014), or weak multivalent binding (Curk, 2016).

#### Simulation of mutations and DFEs

To construct realistic model predictions for mutational returns, we construct a representative gene pool based on the aforementioned landscape fits (see Figure 3A, Figure 3-figure supplement 1A and the Materials and Methods section). The fitness landscapes pool is consistent with the observation that most genes are non-neutral and mostly affect fitness at low expression (Keren et al., 2016), and by design that around 19% of genes are essential (Giaever et al., 2002). We also consider this gene pool in various phases of adaptation, by setting a range of cycle times *T* to generate diversity in background fitness values. We can then also assess whether the coupling of noise and epistasis also depends on background fitness, e.g., whether the simulated mutations follow a diminishing return pattern.

**Figure 3.**
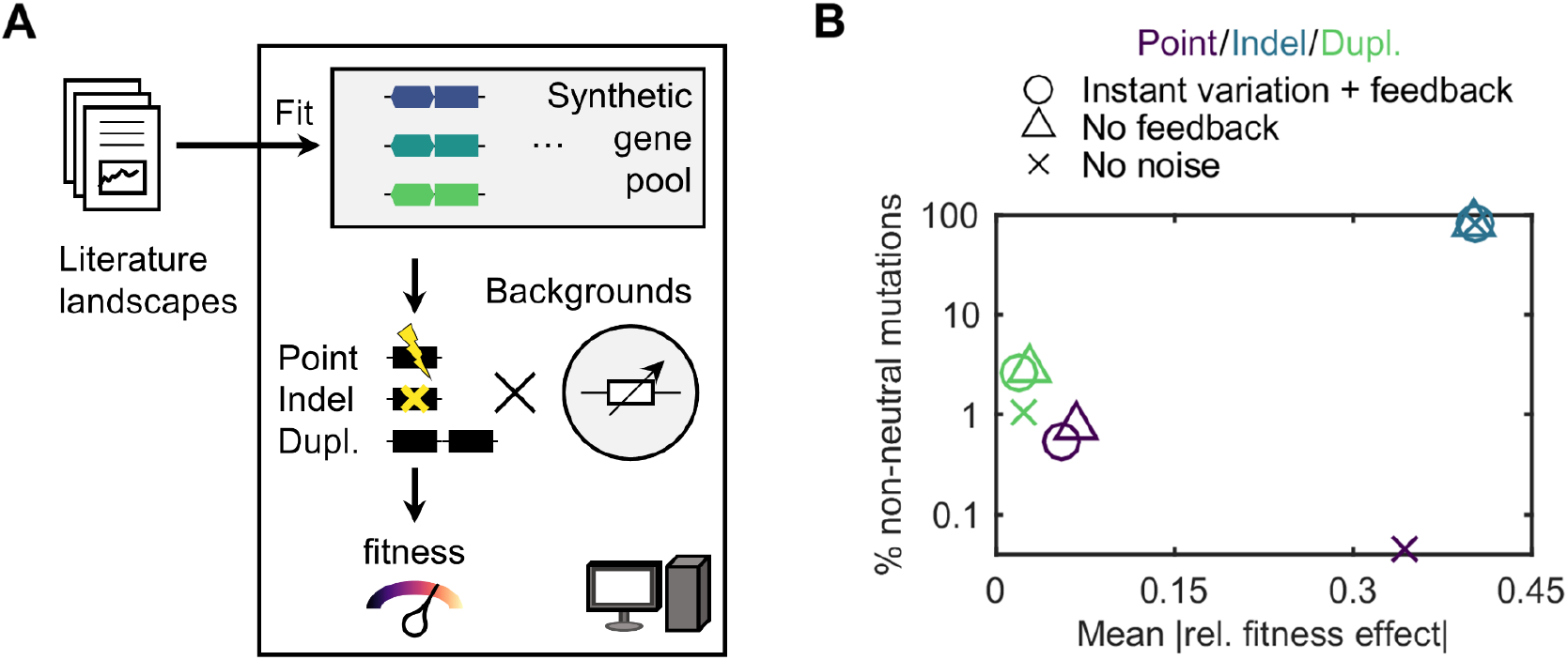
Role of noise on fitness effects of point mutations, indels and duplications. (A) Workflow for generating modelled distribution of fitness effects (DFE) from MEN-model fits of literature fitness landscapes. The fits on landscapes of (Keren et al., 2016) (with expression scaling from (Kulak et al., 2014)) and noise levels from (Chong et al., 2015), fuel construction of a representative gene pool. For one gene at a time, mutations (proxies for point mutations, indels/loss-of-function mutations, and duplications) are imposed while also varying background fitness to mimic different stages in adaptation. This yield simulated DFEs as function of background fitness for various mutation types. (B) Percentage of non-neutral mutations in the DFEs for the aforementioned mutation types. Marker symbols denote the case with instant variation and feedback, only instant variation and no noise.

Given our synthetic gene pool, we consider three noise scenarios: (1) with instant variation and epigenetic feedback, (2) with variation only and (3) completely without noise. These scenarios allow for mechanistic decomposition of the mutational effects. Within these scenarios, we impose three different mutation types to generate distributions of fitness effects (DFEs), which comprise the collection of effects of single mutations in one gene at a time for various values of background fitness. Each type has its own distribution of mutational effects (DME) on gene expression underlying protein X. The mutation types are point mutations, indels/loss-of-function mutations and duplications. The former two are common functional mutations in yeast adaptation (Kryazhimskiy et al., 2014), the latter two are relevant in evolution of many organisms (Murray, 2020; J. Zhang, 2003), such as fungi (Wapinski, Pfeffer, Friedman, & Regev, 2007), *Caenorhabditis elegans* (Farslow et al., 2015), and plants (Panchy, Lehti-Shiu, & Shiu, 2016).

Firstly, we obtain representative point mutations from re-analyzing the data (see Materials and Methods section) on the DMEs from (Hodgins-Davis, Duveau, Walker, & Wittkopp, 2019a, 2019b), where point mutations were randomly chemically induced to shift expression levels of a gene of interest. While based on only 10 lines, the similarity of the DMEs across lines (unimodality, centered around WT expression) suggests this data set is sufficiently representative. The average DME (Figure 3-figure supplement 1C) then allows translation of the landscape as function of expression to a simulated DFE, as function of background fitness. Secondly, the deletions and duplications define an effective expression at zero and twice the original expression. Although indels trivially have the same results with and without noise, we included these in our simulations for an additional validation step (see Simulated DFE comparison to documented diminishing returns), motivating our somewhat arbitrary choice of setting neutrality at ≤0.4% fitness effect.

### Variation generates non-neutrality, feedback recovers most fitness losses for representative mutants

Our earlier theoretical analysis suggested three possible roles of noise: landscape smoothing, extinction delay and, for feedback in particular, recovery of fitness losses. Regarding the first point, we consider the percentage of non-neutral mutations in Figure 3B, which should be larger for smoother landscapes. With noise (circles), the simulations of representative mutations exhibit a large abundance of neutral mutations, 99.5 and 97.4% of point mutations (purple) and duplications (green) respectively. This can be attributed to point mutations causing mainly small expression shifts (see Figure 3-figure supplement 1C) and to toxicity being rare (see Figure 3-figure supplement 1B) respectively. The average mutational effects are also smaller compared to the impactful deletions which are usually non-neutral (see also see Figure 3-figure supplement 1B) but for which noise is trivially irrelevant.

However, without noise (crosses) the percentage of non-neutral mutations plummets about 12-fold and 2.5-fold respectively, yet simultaneously the average mutational effect increases. The loss of instant variation is the predominant cause of this shift, as without feedback (triangles) mutational effects are fairly similar. The instant variation hence generates the noise-driven non-neutrality. Although we theoretically expected more landscape smoothing of the presence of feedback (see Figure 2-figure supplement 2), the aforementioned narrow point mutation DME and rare toxicity mean most non-neutral mutations occur within the relatively narrow part of the landscape, which is smoothened in absence of feedback.

By contrast, we confirm a higher degree of smoothing for essential gene profiles (see Figure 3-figure supplement 2). This translates to a higher percentage of non-neutral mutations for essential genes than for non-essential genes. If we extrapolate the percentage of non-neutral mutations to a higher likelihood of protein X having a genetic interaction affecting expression, this suggests that noise naturally grants essential genes with more interactors. This effect is consistent with the empirical observation that essential genes have about 80% more interactors (see Materials and Methods, (Cherry et al., 2012; Stark et al., 2006)], and that essential genes are more likely to have a hub function (H. Yu, Greenbaum, Lu, Zhu, & Gerstein, 2004). Noise can then be seen as a generator of epistasis.

Expanding on the coupling of noise and epistasis, we bin the mutations per background fitness and determine whether we obtain a positive or negative dependency of the mean fitness effect with background fitness. Surprisingly, the point mutation DFEs show increasing rather than diminishing returns, an effect that is fully attributable to instant variation (see Appendix 1-figure 2). For duplications, the effects are much more subtle, and for indels where noise has no roles, we retrieved the documented diminishing returns with reasonable accuracy.

For the second role of noise regarding extinction, we focus on point mutations, considering the rarity of toxic genes. The non-neutral point mutations can be divided into three situations (see Figure 4A): most of the time (in 84% of the non-neutral mutations), the feedback is not critical for viability, in 14% of the cases the feedback is essential and for 1.7% of these mutations, the population will always decay exponentially. In the latter situation, the feedback notably extends the time to extinction (see Figure 4B), which may permit the time to find a rescuing mutation. On average, feedback increases the number of generations until extinction by 51%, but a broad distribution underlies this number. So, while this situation is rare, feedback is important for these cases.

**Figure 4.**
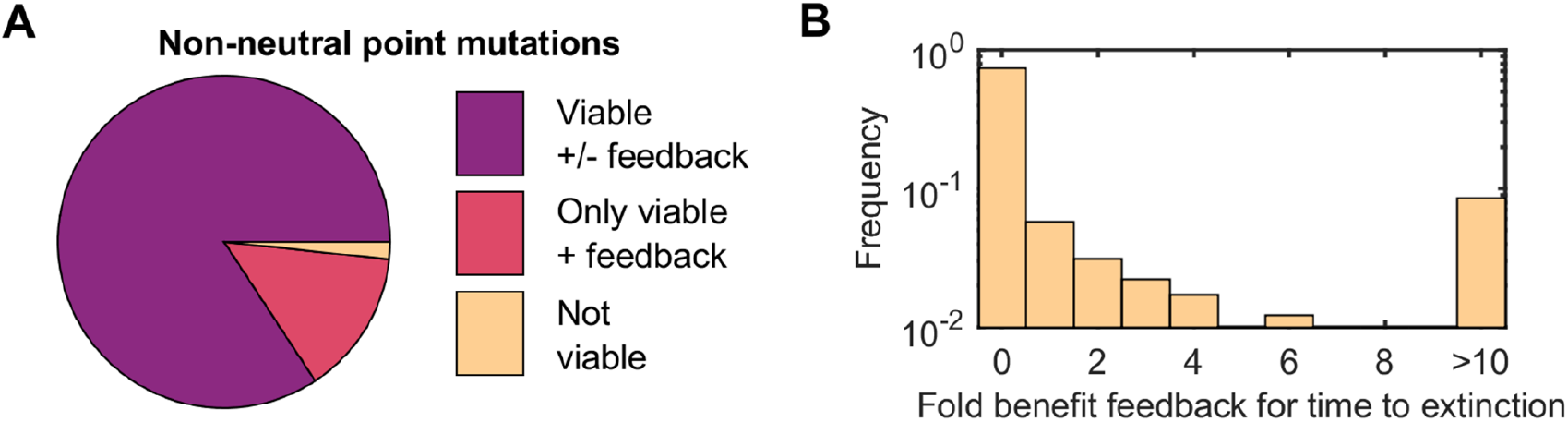
Viability effects of transgenerational feedback. (A) Division of effects of feedback on viability for non-neutral point mutations. (B) Generations to extinction for non-neutral point mutations when the colony is not structurally viable despite feedback.

For the last role, mitigation of fitness losses by feedback, we turn to the vast majority of viable non-neutral point mutations. In Figure 5, heat maps are shown with the magnitude of the fitness penalty that a population suffers when the modelled protein has noise (only instant variation) compared to no noise (vertical axis). As the feedback is never deleterious (see Strict non-negativity of transgenerational feedback effect on fitness), we can plot this against the percentage mitigated when allowing feedback (horizontal axis). Every viable non-neutral point mutation of Figure 4A (red and purple pie) is then placed in a bin along these two axes, with the frequency color-coded (dark blue meaning most abundant).

**Figure 5.**
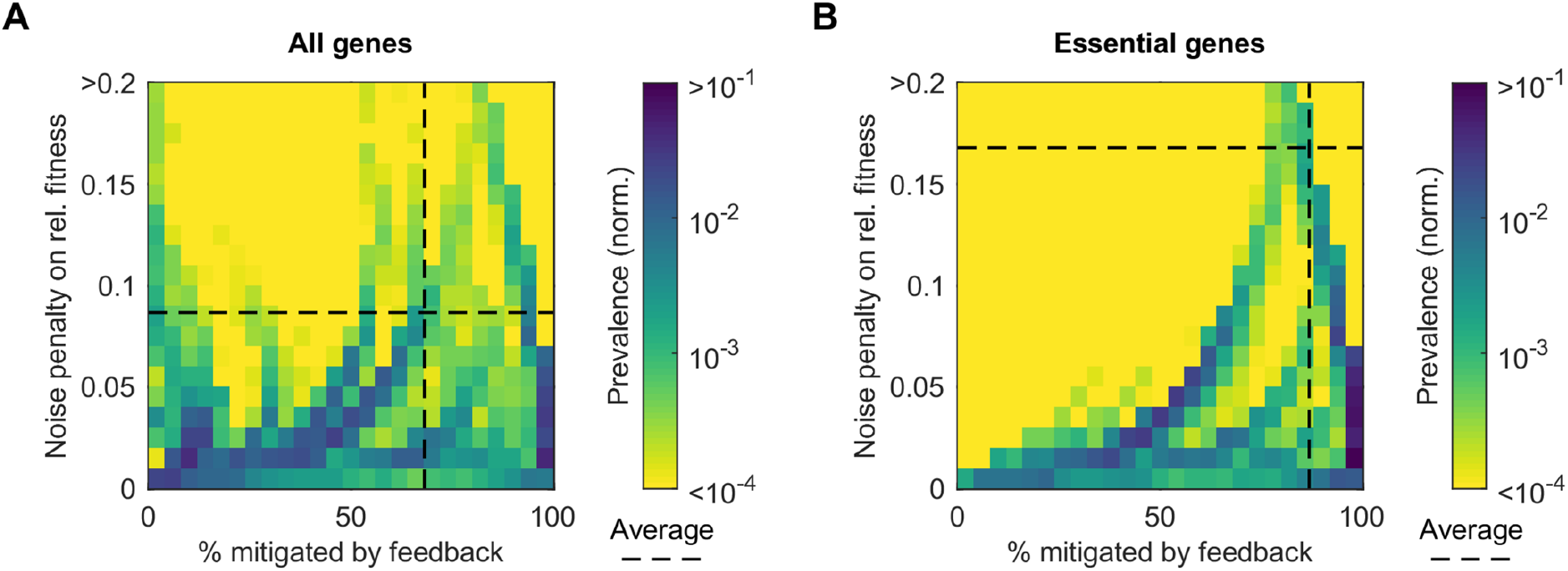
Fitness penalty mitigation by transgenerational feedback. (A) Heat map denoting the frequency (color coded) of non-neutral point mutations with a fitness penalty associated to instant variation by expression noise and a certain mitigation level of feedback. Dashed line denotes average values. (B) The same plot as in (A), only for essential genes.

In Figure 5A, we typically see a 8.7% fitness penalty (dotted line) due to instant variation. Importantly, 68% of this penalty is mitigated by the presence of dosage feedback (a similar conclusion holds for duplications, see Figure 5-figure supplement 1). There seems to be a trend towards a higher percentage of mitigation when the fitness penalty is larger, as can be seen by the locations in the plot of the blue areas from bottom left to top right. This trend is more pronounced for genes for which the highest fitness losses are possible, namely essential genes (Figure 5B). For these genes, the typical fitness penalty is 17%, but the average mitigation level increases to about 87%. This illustrates the contribution of feedback to evolvability; for essential genes in particular, the feedback renders mutations less deleterious than expected, possibly allowing these mutations to become evolutionary accessible.

## Discussion

Epistasis and expression noise are complicating factors for predicting phenotypes and consequentially, the potential course of evolution. To increase the understanding of the mechanical basis of how these factors interact, we have applied a minimal model for epistasis including protein expression noise (MEN-model). Despite the crude approximations of reality, such as binary protein number states, our model accommodated empirical fitness landscapes from (Keren et al., 2016) well, improving on previous fits in 84% of the cases.

Theoretically, several possible roles of noise and the underlying mechanistic basis emerged. Instant variation, the dosage diversity generated intra-generationally, has the potential to smooth the fitness landscape. This landscape smoothing provides for short timescales a causal alternative to the correlation between landscape sharpness and low noise, which has previously been associated to selection for low noise (Keren et al., 2016). The smoothing comes at the cost of a fitness penalty. This penalty is strongly mitigated by epigenetic dosage feedback, a mechanism which acts transgenerationally akin to (Xue & Leibler, 2016) yet adapts to the fitness landscapes as opposed to a fluctuating environment as Xue and Leibler introduced.

To put the feedback in perspective, transgenerational inheritance has been observed at the mRNA level (Houri-Zeevi, Korem Kohanim, Antonova, & Rechavi, 2020), and epigenetic inheritance of doubling times has been linked mechanistically to gene expression noise for the palatinose pathway (Cerulus, New, Pougach, & Verstrepen, 2016). The feedback forms part of a larger panorama of epigenetic inheritance systems (Jablonka & Szathmáry, 1995), but differs in its relative independence of biological requirements. Here, the noise-based transgenerational feedback on protein dosage has an exclusively non-negative effect, also increasing the number of generations until extinction for ill-adapted mean expression levels.

More conceptually, instant variation and feedback combined cause noise to oscillate between the diversity generating role, which has previously been related to bet-hedging (Philippi & Seger, 1989), and another role, partial buffering of deleterious expression levels until a genetic improvement is found. These oscillations resemble an evolutionary LRC-circuit rather than a capacitor as exhibited by other epigenetic mechanisms (Rutherford & Lindquist, 1998). In this LRC-circuit, ‘evolutionary potential energy’ is stored by noise as a capacitor. Upon discharge due to e.g. environmental shock, much of the potential energy is absorbed by transgenerational feedback into the inductor representing noise as a buffer, though at a loss which the feedback cannot fully compensate (the resistance R). After a rescuing mutation, energy returns to the capacitor. It must be noted that in this analogy, the resonance frequency of oscillations is not set by the components but by external, environmental time scales, unlike in electronics.

To determine whether the theoretical expectations hold in realistic circumstances, we constructed distributions of fitness effects (DFE) for non-neutral point mutations, indels and duplications. This was done by simulating a representative pool of synthetic fitness landscapes based on our fits of experimentally determined landscapes in (Keren et al., 2016), and combining these with experimental distributions of mutational effects (DME) on mean protein expression (e.g. (Hodgins-Davis et al., 2019b)). In particular, the rare (~0.5%) non-neutral point mutations best revealed the possible roles of noise.

Firstly, the feedback turns out to play only a small role for smoothing the portion of the fitness landscape that is accessible on the short-term and thus for shaping the target size of non-neutral mutations. Secondly, the feedback has a notable influence on mitigating the deleterious effect of noise, namely reduction of fitness. The feedback recovers most of the reduction the instant variation of protein dosage generates, and is even more important (~87% mitigation) for essential genes. The feedback is therefore realistically pivotal for surviving fitness valleys, particularly for essential genes.

Finally, its diverse roles position noise in important historical evolutionary contexts, see Figure 6. By neutral theory (Kimura, 1983), certain genetic diversity can already be expected by random drift without a significant phenotype penalty, such that there are multiple dark blue dominant genotypes. Moreover, because of bet hedging (Philippi & Seger, 1989), suboptimal genotypes are maintained for more diversity that may become relevant further along adaptation. In addition, feedback mitigates fitness penalties, which allows more genotypes to be maintained than expected. Complementary to this genotypic diversity is the phenotypic diversity generated by instant variation, as light through a divergent lens, which reflects back to the optimal phenotype by the feedback as through a parabolic mirror. The phenotypic variation upon environmental perturbation (Figure 6B) allows time for more permanent, genetic changes to set in (the Baldwin effect (Simpson, 1953)) that mimic the temporarily generated phenotype.

**Figure 6.**
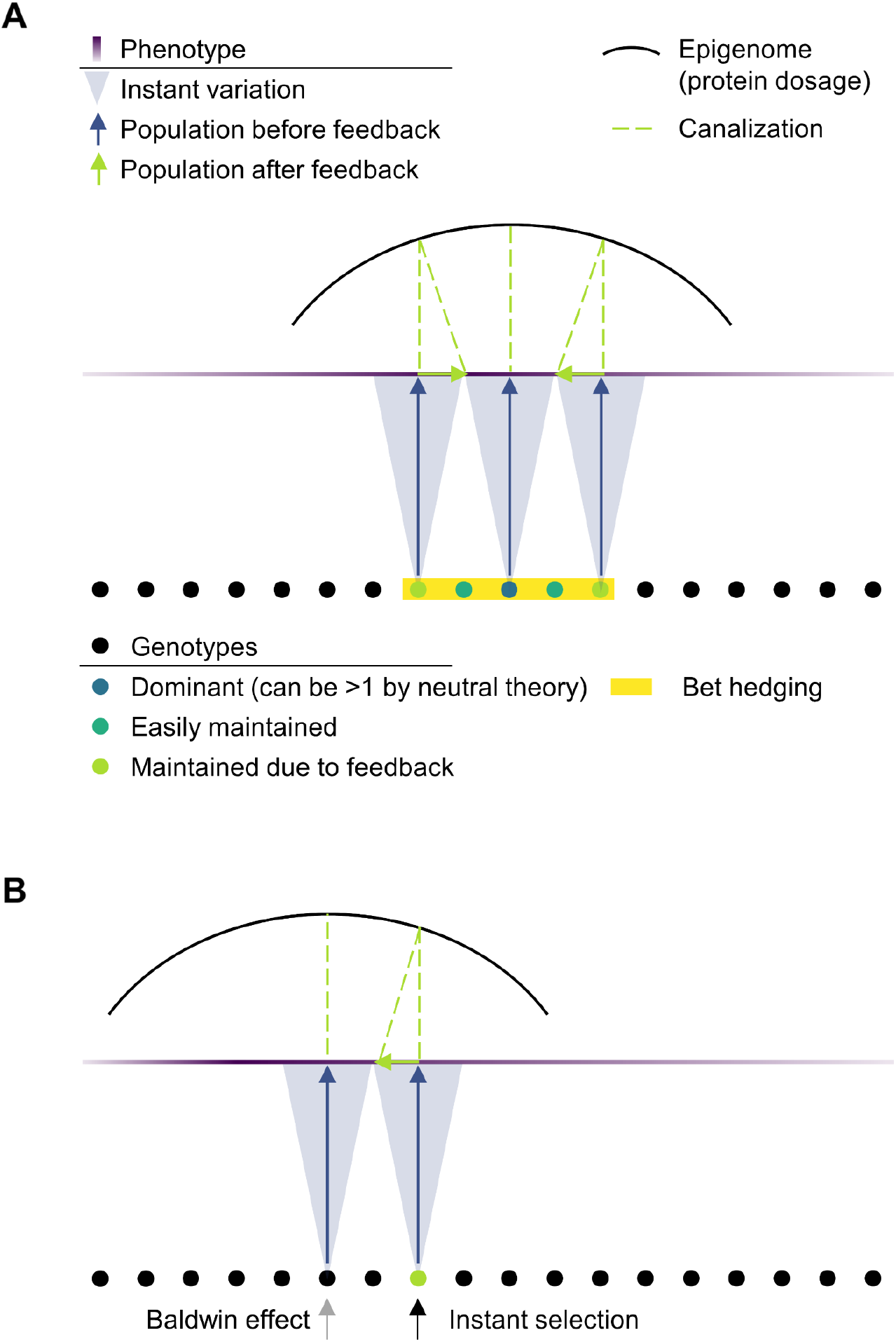
Roles of noise, including transgenerational feedback, in adaptive processes. (A) In addition to the genetic diversity permitted by neutral theory, noise generates phenotypic diversity, for example to prime bet-hedging, during low selective conditions. The instant variation acts as a diverging lens from the genotype plane, which is later refocused onto the phenotype plane through the epigenome acting as a parabolic reflector by making use of transgenerational feedback. This reflection corresponds to a canalization mechanism. (B) After perturbation, the transgenerational feedback as a proper canalization mechanism refocuses the GP-path in the direction of the optimal phenotype. Further genetic adaptation occurs by making use of the enhanced bet-hedging possibilities from panel (A), and if needed the Baldwin effect to consolidate adaptation.

In this view, the continuous reinforcement of the best phenotype through inheritance of acquired copy numbers by transgenerational feedback provides a mechanism for Waddington’s canalization (Waddington, 1942), the driver for genetic assimilation (Loison, 2019; Waddington, 1953). Before the perturbation, the canalization allows for cryptic genotypical diversity and/or lowers the cost of bet-hedging. After the perturbation, the feedback recanalizes the system to the optimum phenotype as the feedback continuously acts on the population. In this way, noise is key in various, complementary evolutionary frameworks.

### Ideas and speculation

Our observations from the simulated DFEs can provide an alternative view on how hub genes emerge. Hubs have many interactors, and often correlate with essentiality (H. Yu et al., 2004), suggesting that strongly connected components become difficult to displace. However, it is possible to invert this causal relation. Due to fitness landscape smoothing by noise, which is theoretically maximal when the fitness drop as function of expression is highest as in essential genes, noise automatically generates many non-neutral expression mutations in essential genes (also in our simulations, see Figure 3-figure supplement 2). It is known that many mutations that affect expression in one gene can actually have their origin in mutations of other genes (Duveau et al., 2021). Therefore, a relatively large non-neutral mutation pool is indicative of a large number of proteins that affect expression of our essential genes, i.e. a large non-neutral pool may involve many interactors. Therefore, essential genes may naturally become hubs by their predisposition for numerous genetic interactions. This provides an unexpected alternative view on how essentiality and connectivity are thought to relate to the evolutionary rate (Fraser, Hirsh, Steinmetz, Scharfe, & Feldman, 2002). In the aforementioned paper the authors propose that connectivity and evolutionary rate correlate by coevolution, and this correlation is not mediated by mutant fitness effects. However, the fitness effect can set the interactivity rather than being the intermediate.

Moreover, if we combine this alternative essentiality-hub causality with the long-term selection for lower noise in these sharp landscapes (Keren et al., 2016), this would translate to a push towards less interactions for essential genes with sharp landscapes. This leads to a reduction of complexity and concordantly, an increase in modularity. The latter is commonplace in nature with multiple theorized origins (Wagner, Pavlicev, & Cheverud, 2007), and selection on noise can be one more source. Therefore, another role for noise may surface through the linkage of noise and genetic architecture.

## Materials and methods

### MEN-model fits on fitness landscapes from literature

We considered the fluorescence values for different promoters and original fits from (Keren et al., 2016) in glucose conditions. We related observed fluorescence to known WT dosage by dividing all fluorescence data from the synthetic promoters by the median ratio of fluorescence under endogenous expression and copy numbers from (Kulak et al., 2014). This was only possible for 73 of the 81 landscapes. Data at zero expression was supplemented by an essential gene data set, consisting of the null mutants in yeast strain background S288c from SGD (Cherry et al., 2012) (date of access 01-08-2019).

Fitting model values of λ_*max*_/2 (with λ_*max*_ being our population growth factor per generation in Equation 3) to the observed landscapes was performed in Matlab R2016a (as are all calculations throughout this paper) using the native *fminsearch* to minimize the sum of squared residuals. For essential genes, the fitness according to the definition of (Keren et al., 2016) at zero expression was set at 0, although any value below 1 would represent an essential gene here. However, setting the fitness at 0 works well to correctly match modelled (non-)essentiality and actual (non-)essentiality 92% of the time. For the calculation of the adjusted R-squared of the fits of (Keren et al., 2016), essentiality data is excluded, as it was not considered in that paper.

Fitting parameters *c*, *k* and *d* and *V* were restricted to 0 to ∞, −10 to 10, 0 to 2 and 0.1 to 1 (the latter as feasible range in (Chong et al., 2015)) respectively. We note that this *V* only has relevance in the context of these fits, as it represents an effective noise originating from the various synthetic promoters combined. Fits are then further fine-tuned with the Matlab’s R2016a *fit* function, where the restrictions on *k* are fully relaxed to also provide *k* with a (67%) confidence interval.

### Simulation of representative DFEs

To generate the DFE as a function of background fitness, we first create a representative gene pool based on the model fits from the 61 fitness landscapes of (Keren et al., 2016) whose fits quality improved the original fits from that paper. Within the class of essential and non-essential genes separately, we resample new progeny shapes from random combinations of fitted model parameters *c*, *k* and *d*. As can be inferred from Appendix 1-figure 1C, essential genes (with large effect of deletions) constitute a relatively large part of the data set and we correct for this bias. This yields a balance between various fitness landscapes at shown in Figure 3-figure supplement 1B such that approximately 19% (Giaever et al., 2002) of all combinations corresponds to essential gene profiles (5000 simulated genes in total except for Appendix 1-figure 3). We also assume fitness at WT expression is at >95% of the maximal value, such that WT is relatively optimized, and then normalize all fitness values for that gene to the value at WT expression. By setting different values for cycle time *T* (between −50% and ~+5% relative to WT) and randomly assigning a noise level drawn from (Chong et al., 2015), we generate simulated fitness landscapes as function of expression, where background fitness is then defined as the fitness at WT expression at the various cycle times. For Figure 3, Figure 3-figure supplement 2 and Figure 5-figure supplement 1B, the absolute mutational fitness effects considered are compared to the background fitness of each respective scenario (instant variation + feedback, only feedback or no noise), otherwise background fitness with instant variation and feedback is assumed.

To convert these landscapes to DFEs, we transform for point mutations the expression axis to mutation frequency using the DME data from (Hodgins-Davis et al., 2019a, 2019b). After converting the observed counts to a density by normal kernel density smoothing, we interpret the control distributions as point-spread functions blurring the real mutation distributions. We then retrieve the latter by Lucy-Richardson deconvolution (Fish, Brinicombe, Pike, & Walker, 1995), as applied in Matlab’s *deconvlucy* of the observed mutation distributions. Using 201 uniform samples of the average DME, we obtain the DFE by expression mutants, as a function of background fitness. DMEs of the indels and duplications are trivially only non-zero at zero and twice the WT expression.

We then consider the DFEs for the three scenario’s: with/without noise (instant variation + feedback) where fitness follows from equations 3 and SI.8, and with feedback suppressed (SI.12). To avoid singularities in the calculation of the cumulative distribution function, we set zero and infinite noise values to 10^-20^ and 10^20^ respectively instead.

### Essentiality and interactions

Essential gene data is from the SGD Project (Cherry et al., 2012), date of access 1 August 2019, interactions data from BioGRID (Stark et al., 2006), date of access 3 February 2022. As our interactions of interest are genetic, we consider the number of interacting proteins per protein of interest for which a genetic but not a physical interaction exists. The essential genes are those genes that yield inviability with a null mutations in a S288C background.

## Acknowledgements

We thank Marieke Glazenburg and Djordje Bajić for careful reading of the manuscript. LL gratefully acknowledges funding from the European Research Council (ERC) under the European Union’s Horizon 2020 research and innovation programme (Grant agreement No. [758132]).

## Figures

**Figure 1-figure supplement 1.**
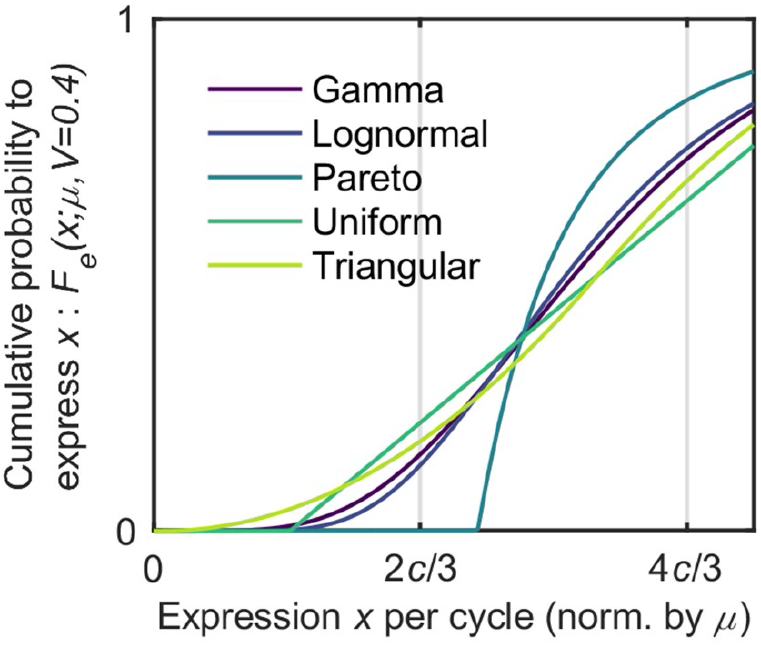
Five cumulative distribution functions for fixed *V* (at 0.4), as function of expression per cycle, rescaled to the mean expression. Horizontal axis ticks indicate the low and high states in the MEN-model.

**Figure 2-figure supplement 1.**
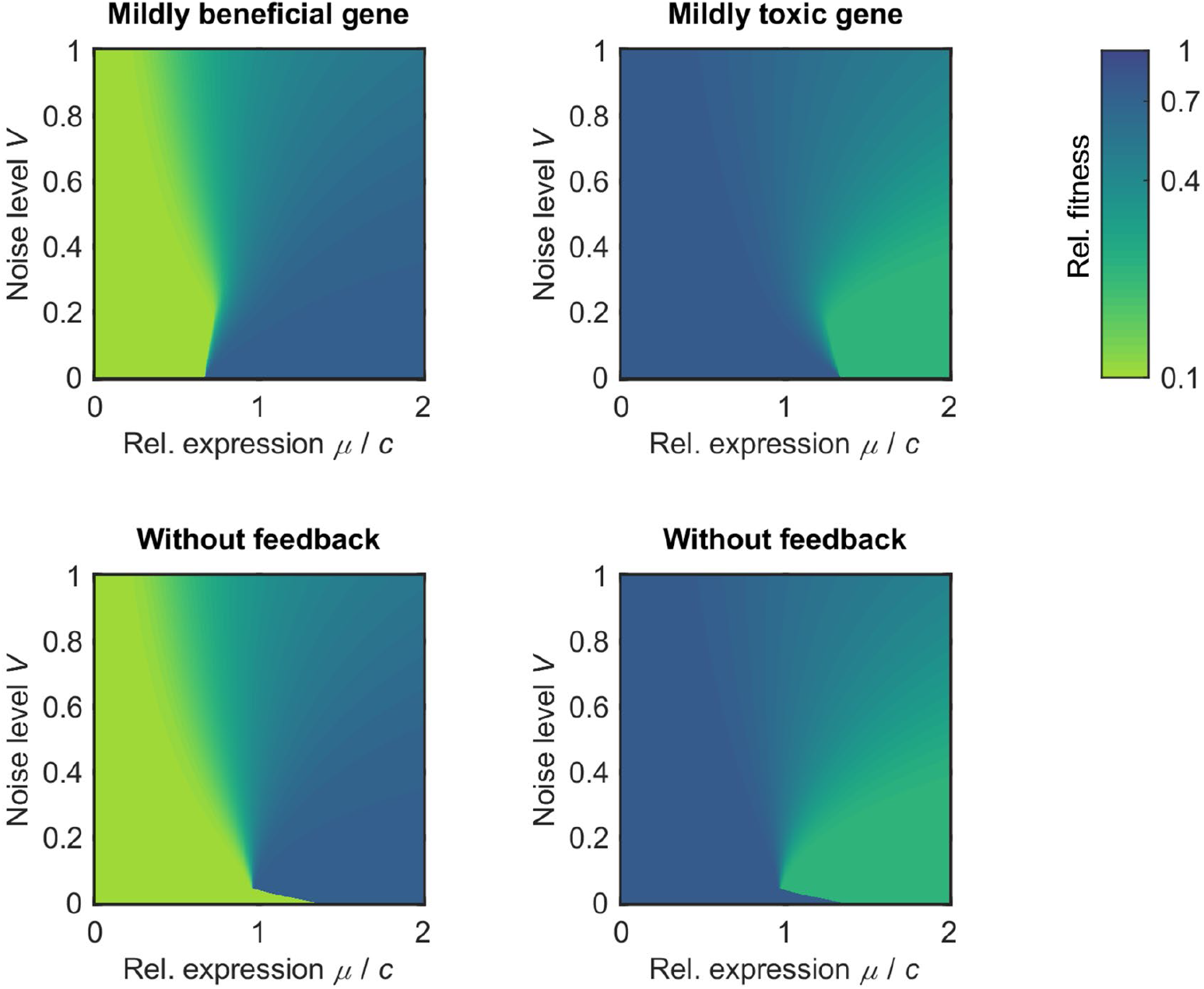
Two fitness landscape types with structurally viable regimes, with and without feedback. Landscapes are given as heat maps plots as function of relative expression *μ* (scaled by tipping point concentration *c*) and noise level *V* (from 0 to 1, which covers 99.8% of the genes in (Chong et al., 2015)). Blue to light green colors denote the log of relative fitness *ω_r_* All plots assume a gamma cdf *F_e_*, |*k*|=3 and *d*=0.8.

**Figure 2-figure supplement 2.**
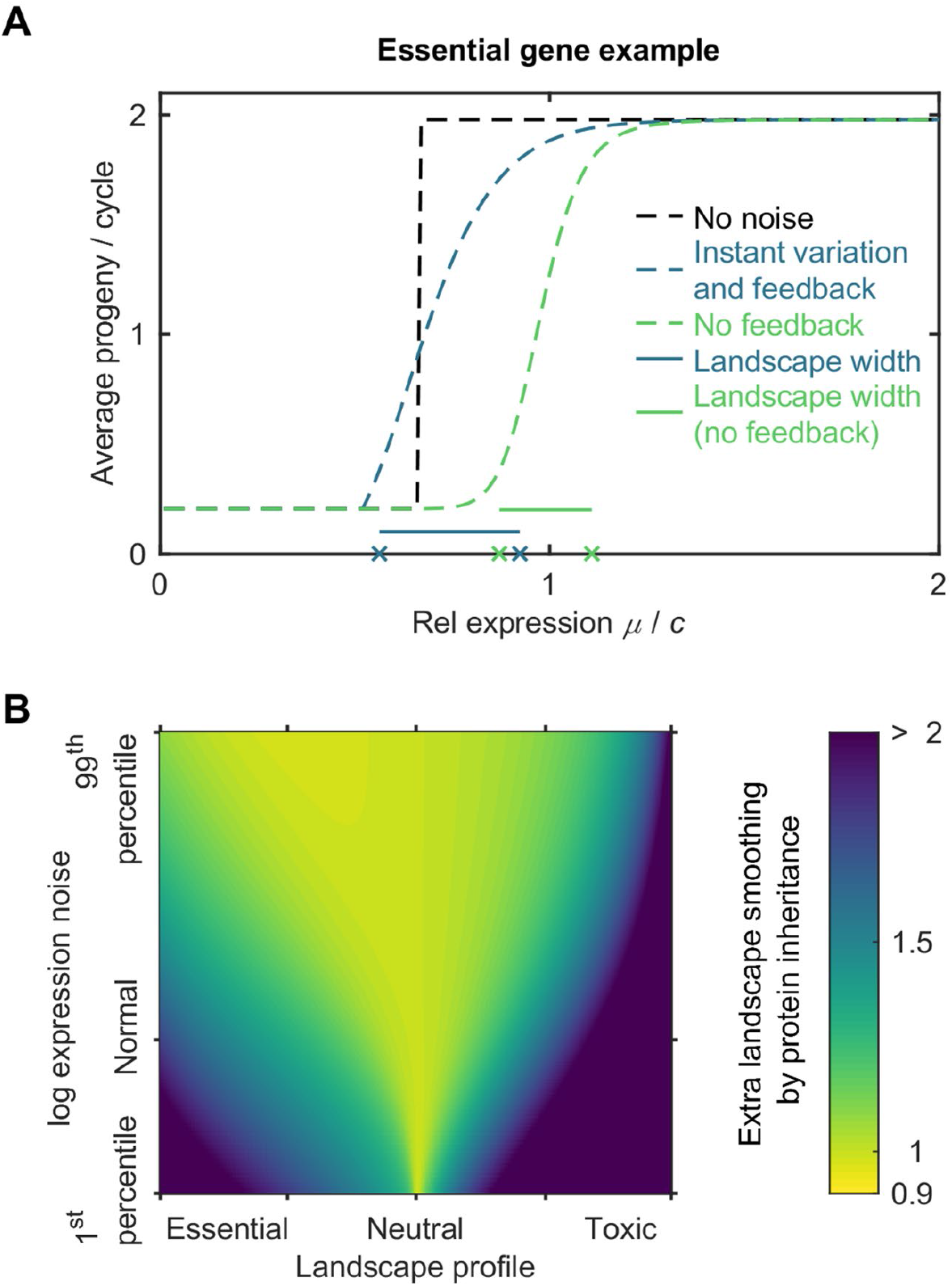
Theoretical effect of transgenerational feedback on smoothing of fitness landscapes in the MEN-model. (A) Example fitness landscape to illustrate the definition of landscape width as expression range for which fitness falls between 10% and 90% of the maximum for a given noise level *V* and progeny difference between the high and low state. (B) Ratio of widths between the cases with and without feedback. We assume a gamma cdf *F_e_*,, and | *k*|=15, a typical fitted value fitted (see Appendix 1-figure 1C).

**Figure 3-figure supplement 1.**
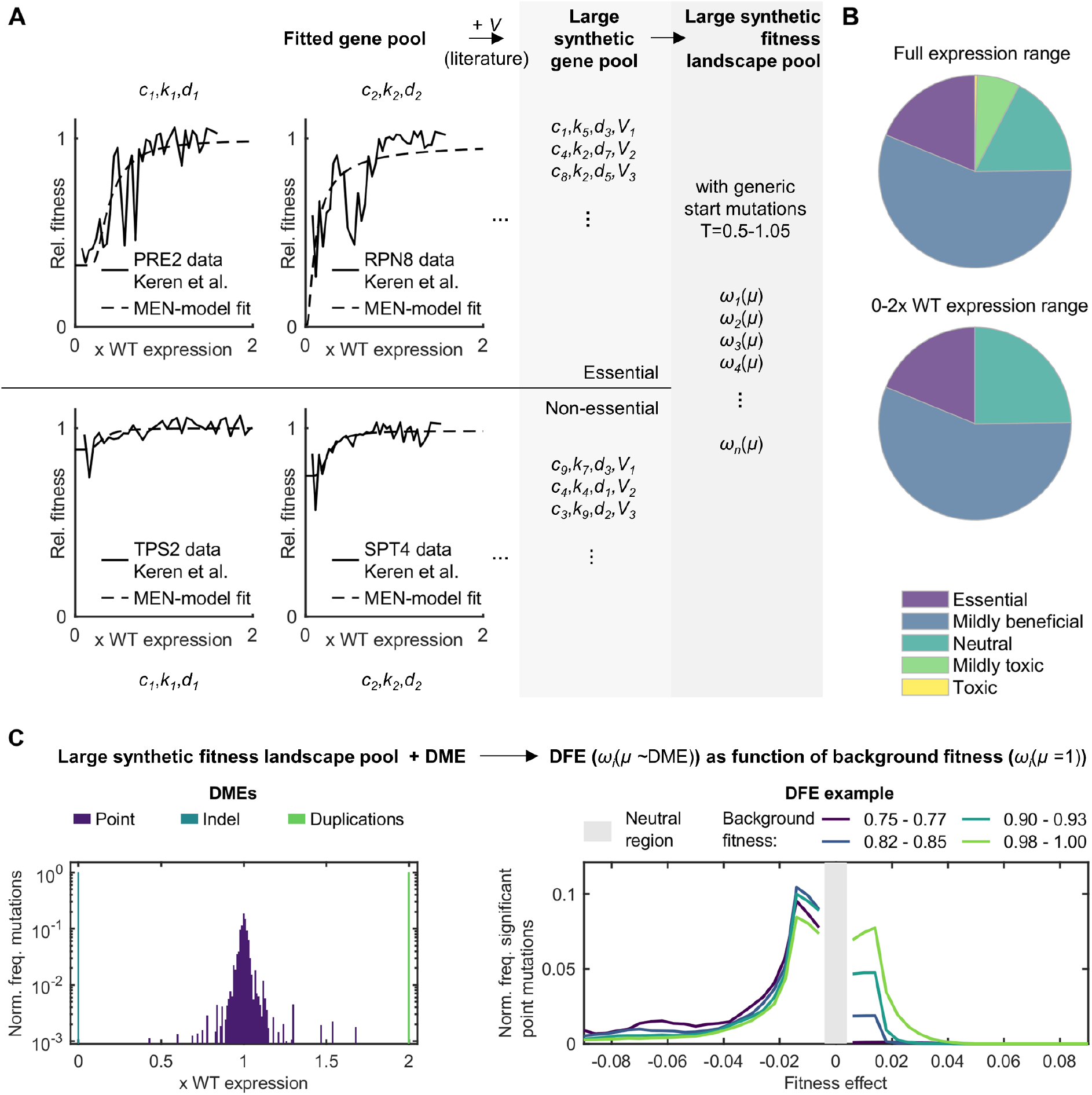
Workflow for generating modelled distribution of fitness effects (DFE) from MEN-model fits of literature fitness landscapes.(A) Construction of the large simulated gene pool with corresponding fitness landscapes, based on MEN-model fits of landscapes of (Keren et al., 2016) (with expression scaling from (Kulak et al., 2014)) and noise levels from (Chong et al., 2015). Displayed empirical fitness landscape examples are selected for good coverage across 0 to 2 WT expression levels for instructive purposes. This panel uses the mean progeny per cycle as fitness definition as in (Keren et al., 2016) instead of Equation 3. (B) Fractions of simulated fitness landscapes in the categories: essential (zero fitness at zero expression), mildly beneficial (viable across expression range, but better at high expression, with fitness rise > 0.05), neutral (maximum fitness differences across expression range < 0.05), mildly toxic (viable across expression range, but better at low expression, with fitness rise > 0.05) and toxic (inviable at high expression). (C) Construction of the DFEs as function of background fitness using the distribution of mutational effects on expression (DME) for promoter mutations re-analyzed from (Hodgins-Davis et al., 2019a, 2019b). Neutral (fitness effect < 4e-3 as a proxy for non-significance based on the experimental resolution in (Johnson, Martsul, Kryazhimskiy, & Desai, 2019)) and non-viable mutations are removed from the DFE, which is smoothened only here for visualization. Background fitness values are binned, ranging from 0.75 to 1.05.

**Figure 3-figure supplement 2.**
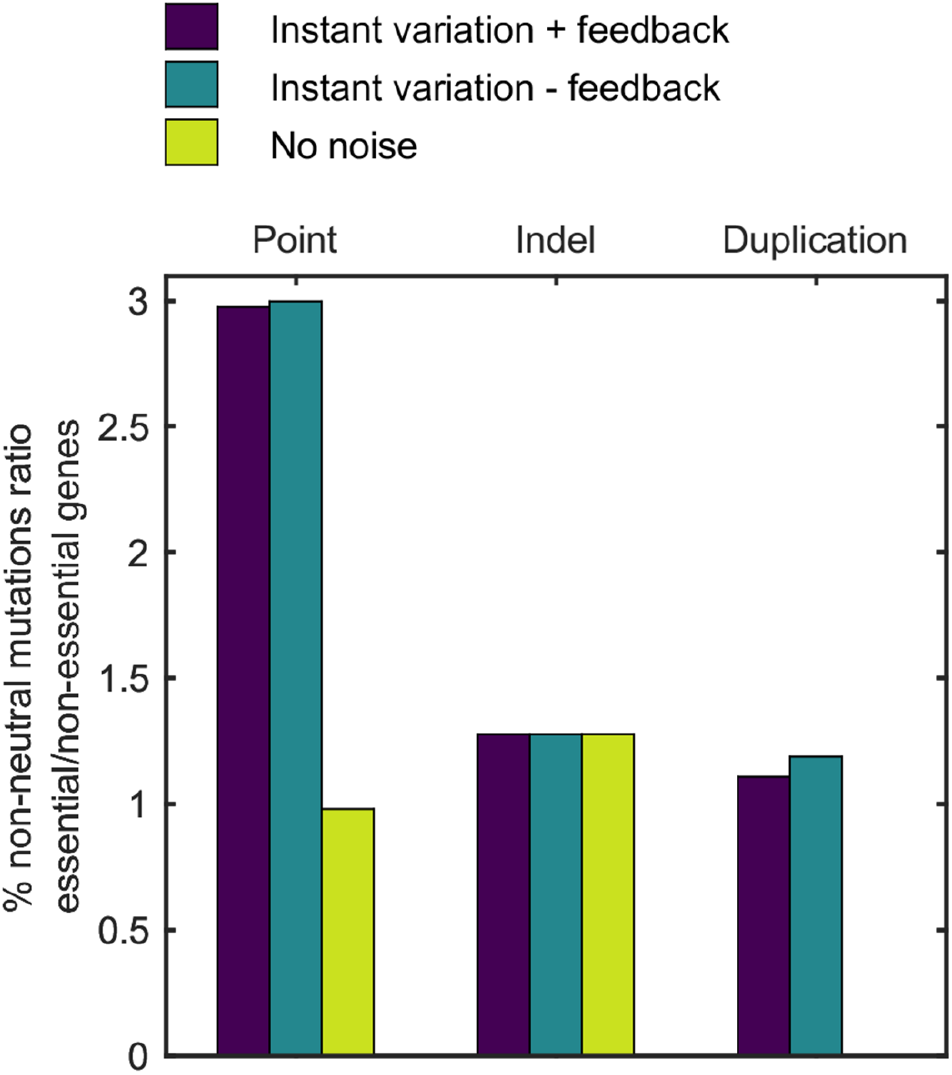
Influence of essentiality on mutational effects. Bars represent the ratio between the percentage of non-neutral mutations in essential and non-essential genes, for simulated point mutations, indels and duplications as in Figure 3.

**Figure 5-figure supplement 1.**
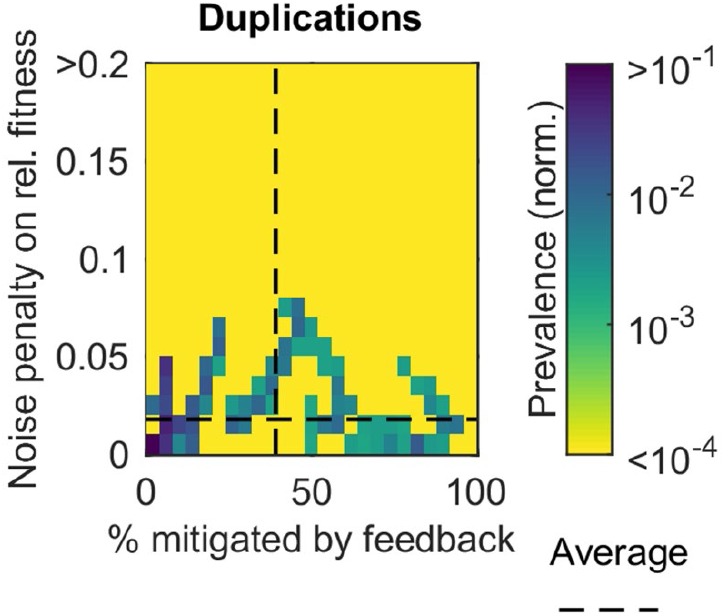
Effects of noise and feedback on mutational returns of duplications. The heat map denotes the frequency (color coded) of non-neutral duplications with a fitness penalty associated to instant variation by expression noise and a certain mitigation level of feedback. Dashed line denotes average values. The same duplication DFE is considered as in Figure 3.

---

**Appendix 1-figure 1.**
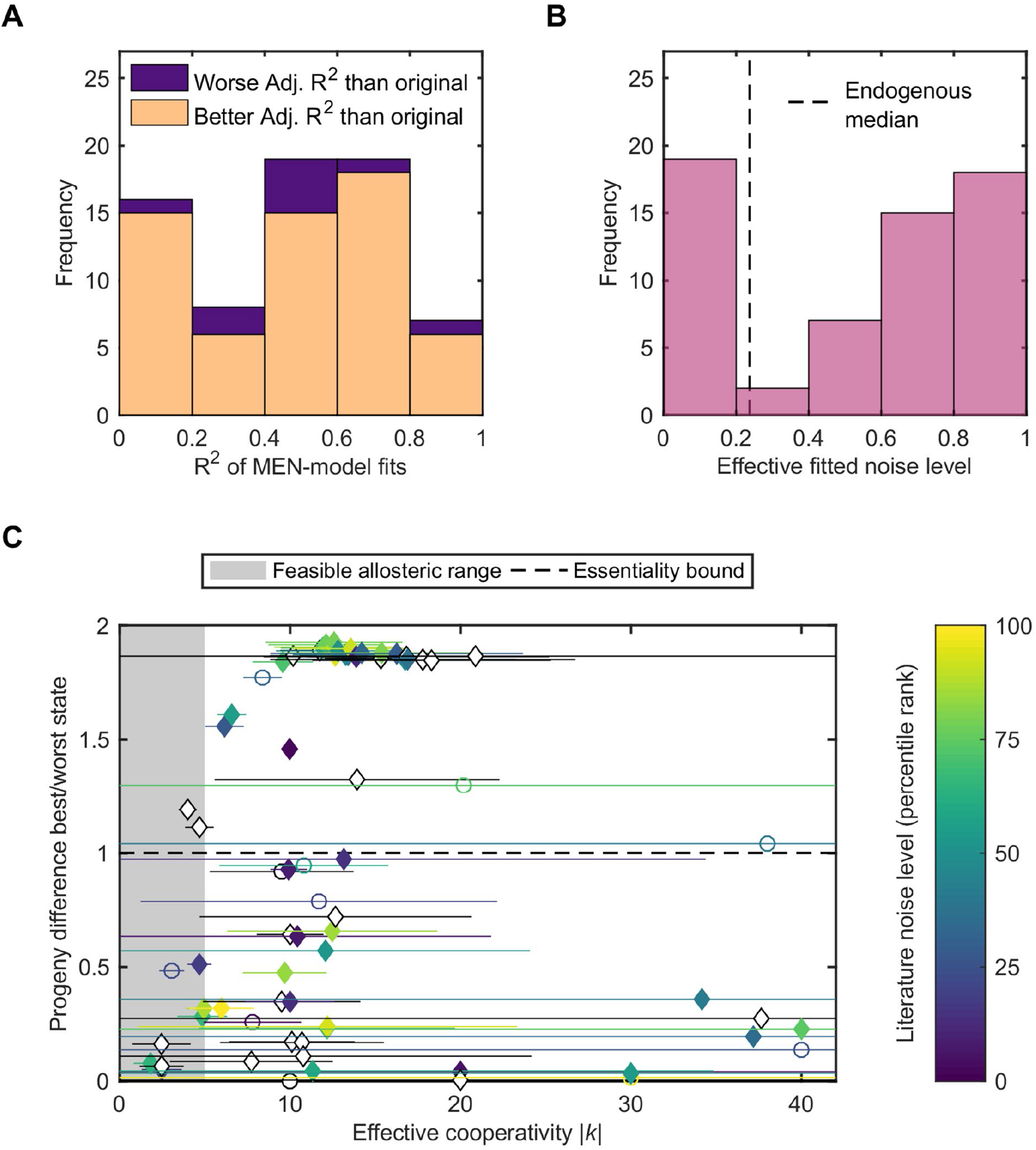
MEN-model validation and assessment of fitted fitness landscapes.(A) Goodness of fit of the MEN-model compared to the original fits of (Keren et al., 2016) on the associated fitness landscapes. (B) MEN=model fit estimate of the effective noise of all promoters combined from the landscapes for which the MEN-model improves the fits and noise level (coefficient of variation) estimates of (Chong et al., 2015) are available. (C) Fitness drop between states plotted against the parameter *k*, color coded with the literature noise values (Chong et al., 2015) (converted to ranks). No color for marker filling is given when for this gene no noise estimate was available (then only black edges and error bars, 67% confidence intervals) or when the model fit is worse than the original.

**Appendix 1-figure 2.**
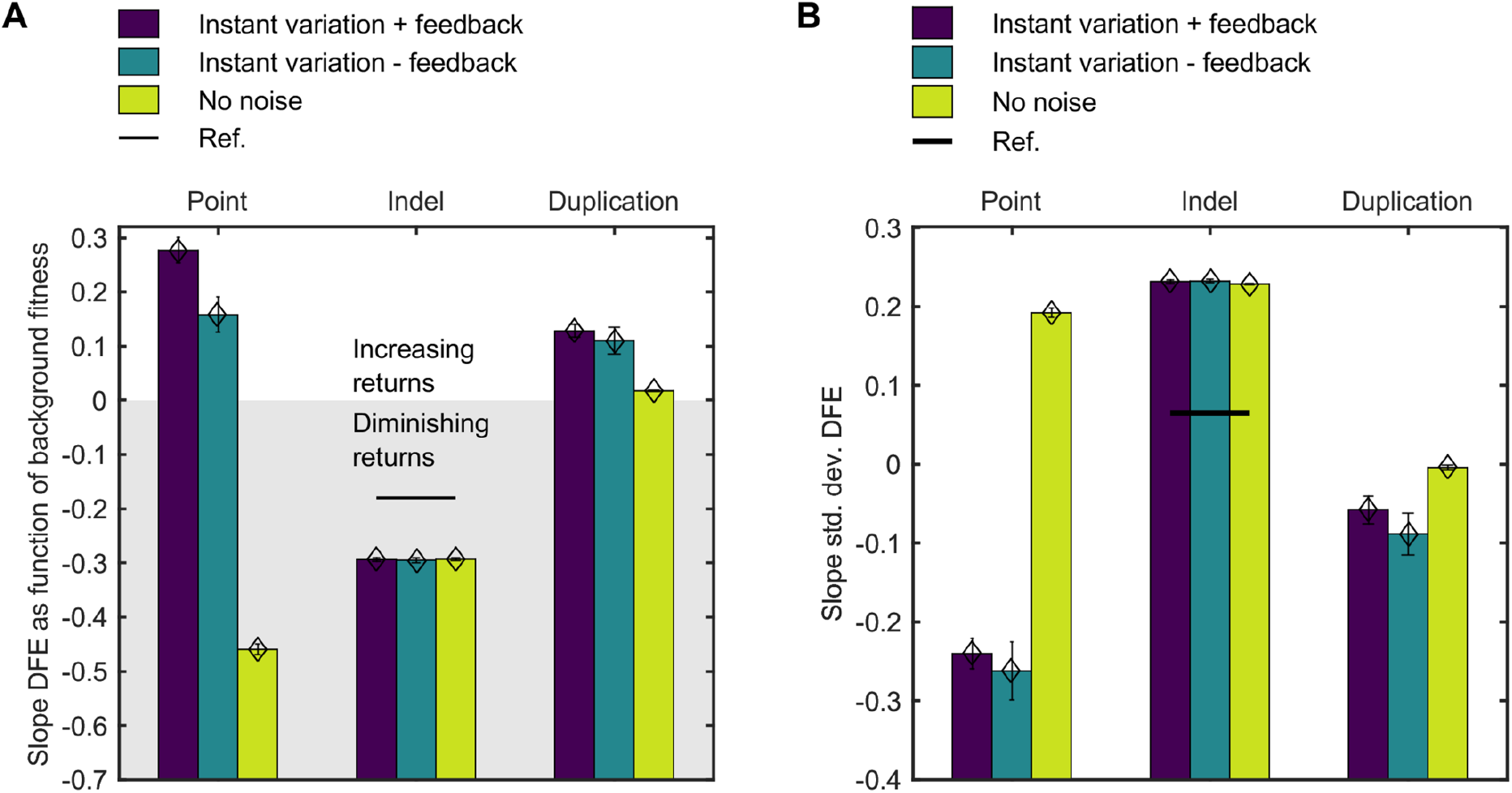
Effects on noise on the mean and standard deviation of the DFE. Fits are performed on the standard deviations of simulated DFEs as function of background fitness, with instant variation and feedback, instant variation alone and without noise. Positive slopes represent increasing returns (A) or noisier (B) returns as background fitness increases. Fits follow from weighted least squares (WLS) on the mean or standard deviation bootstrapped per 0.025 background fitness bin, with weights as the reciprocal of the variance of the bootstrap values. DFEs and DMEs considered are for the point mutations, indels and duplications as in Figure 3, only disregarding lethal mutations for more appropriate comparison to the experimental reference value from (Johnson et al., 2019), relevant for indel DFEs.

**Appendix 1-figure 3.**
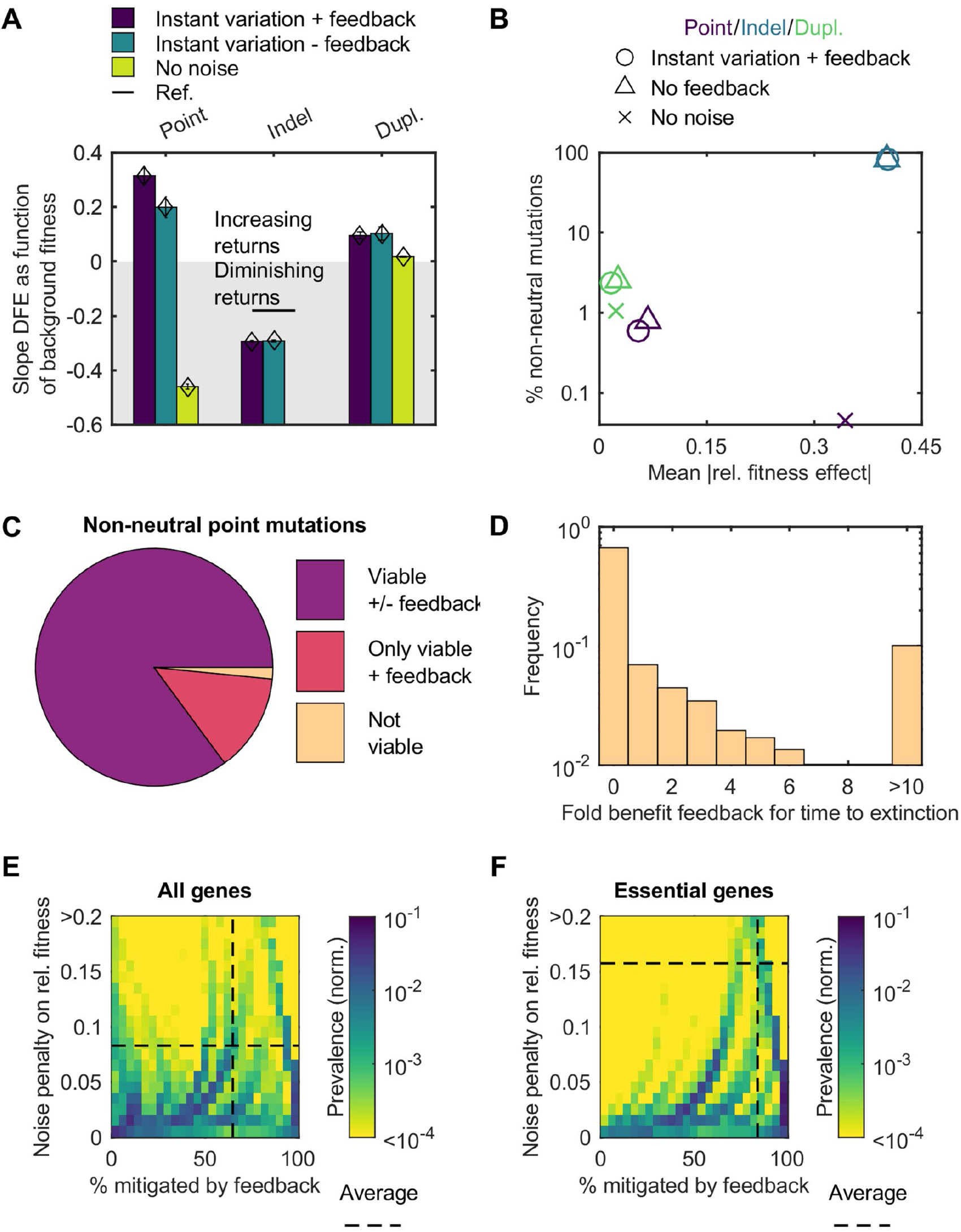
Effects of noise on mutational returns when assuming linear scaling between log noise level and log expression (log *V* / log *μ* = −0.27) mimicking observations in the low expression regime from (Keren et al., 2015). Mutational pool is 500 instead of 5000 genes, but otherwise no changes are made in the workflow compared to Appendix 1-figure 2A (for panel A) and Figure 3 (other panels). (A) Fits on the mean of simulated DFEs as function of background fitness, with instant variation alone, with instant variation and feedback, and without noise. Negative mean slopes represent diminishing returns. Fits follow from weighted least squares (WLS) on the mean bootstrapped per 0.025 background fitness bin, with weights as the reciprocal of the variance of the bootstrap values. (B) Percentage of non-neutral mutations in the DFEs for the DMEs considered. Marker symbols denote the case with instant variation and feedback, only instant variation and no noise. (C) Division of effects of feedback on viability for non-neutral point mutations. (D) Generations to extinction for non-neutral point mutations when the colony is not structurally viable despite feedback. (E) Heat map denoting he frequency (color coded) of non-neutral point mutations with a fitness penalty associated to instant variation by expression noise and a certain mitigation level of feedback. Dashed line denotes average values. (F) The same plot as in (E), only for essential genes.

## Appendix

### Model implementation details

For our model, we assume a constant size for all cells, and hence no dilution. We turned to data from (Chong et al., 2015), where many proteins in yeast were tagged with GFP such that the protein distribution across the population could be determined. By examining the coefficients of variation, which we define as noise level *V*, we get an indication on whether the noise levels on gene products are gene specific or have common noise sources, such as dilution. The floor value of *V*, which links to the common sources is 0.1, while the typical *V* (median) is 0.2. Using a crude root-mean-square decomposition of *V* suggests the common sources as dilution are typically only responsible for 25% of the noise, so idiosyncratic noise contributions dominate.

Additionally, we assume no degradation of proteins. To see how stringent this assumption is, we assessed the degradation in several model systems. In *E. coli*, there is no noticeable degradation for 93 to 98% of the proteome (Nagar et al., 2021). In *Schizosaccharomyces pombe* and *S. cerevisiae*, degradation is not a factor for protein abundancy for around 85% of the proteins (Christiano, Nagaraj, Fröhlich, & Walther, 2014). Finally, we note that for *C. elegans*, the generation time for the multicellular organism is on the scale of the typical protein half-life (Dhondt et al., 2017; Muschiol, Schroeder, & Traunspurger, 2009). This suggests that protein degradation is important for roughly half the proteins. In conclusion, explicitly ignoring degradation is inconsequential for the standard microbes but may become a factor for higher order organisms.

Furthermore, we simplify protein states to two discrete states to maximize analytical tractability. The rationale is that the effect of stochasticity in the model is already incorporated as soon as there is more than one state. Moreover, as shown further on most empirical fitness landscapes of (Keren et al., 2016) are consistent with one or two protein states with distinct progeny levels (see Appendix 1-figure 1A). The natural concentration scale that we can use to define the states is the tipping point concentration *c* in the Hill progeny curve from Equation 1. Ideally, we define the state symmetrically around *c* and as equally spaces as possible. However, the halving of the concentrations upon generating the progeny and the need to revert to the same states every cycle limits our choice of binning. As a compromise, we set our high and low state to *X=2c/3* and *X=0* respectively before protein production and to *X=4c/3* and *X=2c/3* respectively after production. The relevant progeny, after production, is then the Hill curve from Equation 1 evaluated at *X=4c/3* and *X=2c/3*, so for the high and low state *gh* and *gl* respectively:

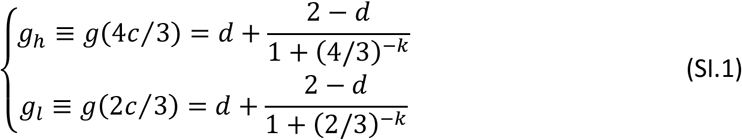

Sometimes, the progeny-protein scaling assumption can be justified from the bottom-up, such as in the case of polarity establishment in *S. cerevisiae*. There, polarity success, which is essential for cell division, depends in an almost binary fashion on Cdc42p concentration (Brauns et al., 2020). In such a case, the progeny function also classifies as a mesotype (Daalman, Sweep, & Laan, 2021).

Due to the stochastic protein production of X, a cell can switch between protein occupancy state with every cycle. We define the cumulative distribution function (cdf) for random variable *x*, the added protein per cycle, as *F*_e_(x; *μ, V*), with *μ* as the average production per cycle and *V* as the coefficient of variation, or noise level, of the protein production per cycle. We only consider the probability values of the two state transitions from the high state (before production) to the low state or low state to low state, which can be defined as *F_l_* and *F_h_* respectively. This is because the other two transitions simply follow from noting the probability to end in any state starting from the high (or low) state is trivially 1. To end in the low state from the high state (at *X=2c/3*), the production x may not exceed 2c/3, otherwise the total ends in the high bin again (at *X=4c/3*). Similarly, to end in the low state from the low state (at *X=0*), the production x may not exceed 4c/3, otherwise the total ends in the high bin again (at *X=4c/3*). Therefore, the probabilities *F_h_* and *F_l_* are given by:

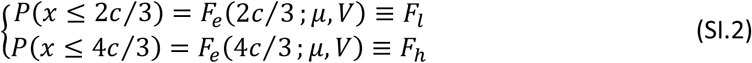

where the mean μ depends implicitly on the cycle time; if the cycle time is longer, the mean increases and the probabilities to reach the high state after production are higher.

Concretely, the precise values of *F_h_* and *F_l_* depend on the choice of the distribution for the cdf. Figure 1-figure supplement 1 shows the cdfs for five different choices, including the gamma cdf, which is a sensible choice for modeling the production. A gamma distribution follows from adding exponential random variables, the latter being suitable to model protein numbers from expression bursts (see (Friedman, Cai, & Xie, 2006), motivated therein by experiments in references (Cai, Friedman, & Xie, 2006; J. Yu, Xiao, Ren, Lao, & Xie, 2006)). In short, we see that fixing the first two moments of the distribution is rather restrictive for the precise values of *F_h_* and *F_l_*, even for unbiological choices for the protein production cdf (such as a triangular cdf), except for the Pareto distribution. Therefore, our model results are not sensitive to the choice of cdf, unless the real distribution of the protein expression bursts are very far from expectation.

### MEN-model fitness comparison with literature

After *n* generations, the number of cells in each protein state is given by:

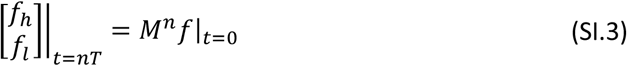

If we decompose the initial state *f*|_*t*=0_ into the eigenvectors *v*_1_ and *v*_2_ of *M* (with appropriate weights *a_1_* and *a_2_*, and *λ_1_* and *λ_2_* as the respective eigenvalues with *λ_1_* ≥ *λ_2_*), then we have:

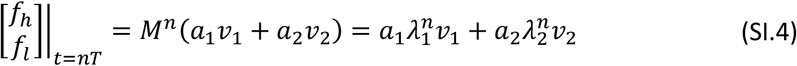

After sufficient generations, only the term with the largest eigenvalue remains, so:

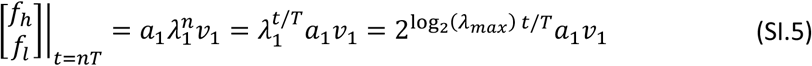

From this expression, we note the time to double the state occupancy is *T*/log_2_(*λ_max_*), so the fitness, which is the reciprocal of this time, becomes *ω*=log_2_(*λ_max_*)/*T*, the expression in equation 3.

We can write the eigenvalues, of which we need the largest one for equation 3, as:

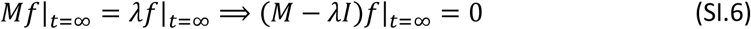

This is routinely solved by setting det(*M* – *ΛI*) = 0, which for a 2×2 system reduces to:

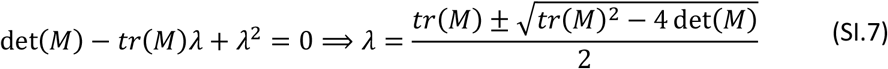

The determinant can be written as:

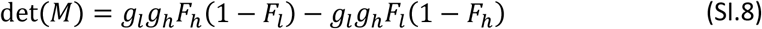

Generally, the first term is larger than the second, as *F_h_* > *F_l_* and thus also 1 – *F_l_* > 1 – *F_h_*. Assuming relatively low noise levels (e.g. those found in *S. cerevisiae* where the median value is 0.2 (Chong et al., 2015)), we can state *F_h_* ≫ *F_l_* and 1 – *F_l_* ≫ 1 – *F_h_*, and then we can approximate the determinant as

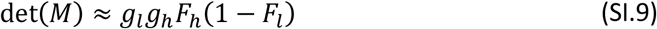

The largest eigenvalue is then:

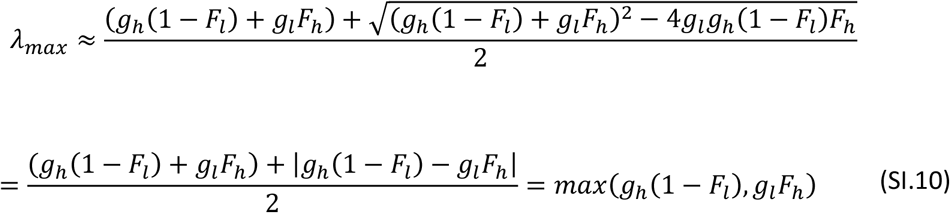

Combining this result with equation 3 yields equation SI.11, provided that there is sustainable growth to define fitness *(ω_r_* > 0):

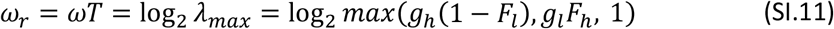

Interestingly, we note a corollary with the fit function in (Keren et al., 2016), which the authors employed to fit their empirical fitness landscapes. Their fit function consists of a product of two sigmoids (from (Chechik et al., 2008)), fitting a fitness which they defined as the number of progeny compared to WT, equating to *λ_max_*/2 in our model. We see in our expression for *λ_max_* the contours of the product of two sigmoids, *g* and *F*, only this time motivated from the bottom-up, and incidentally, with three free parameters less (4 instead of 7). While the double sigmoid worked well for authors of (Keren et al., 2016) on their fitness landscapes, our model construction provides the insight why their fit function worked so well.

### Strict non-negativity of transgenerational feedback effect on fitness

Given suppression of feedback effectively resets the state vector *f* to the same value at every iteration, the state equation 2 can be modified for the absence of transgenerational feedback as:

The state equation for the case when feedback is suppressed can be written as:

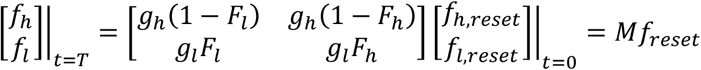

We define the *f_reset_* as the state vector in equilibrium when there is no selection, so when *g_h_*=*g_l_*=2. In that case, *λ_max_*=2 as all cells produce two daughter cells:

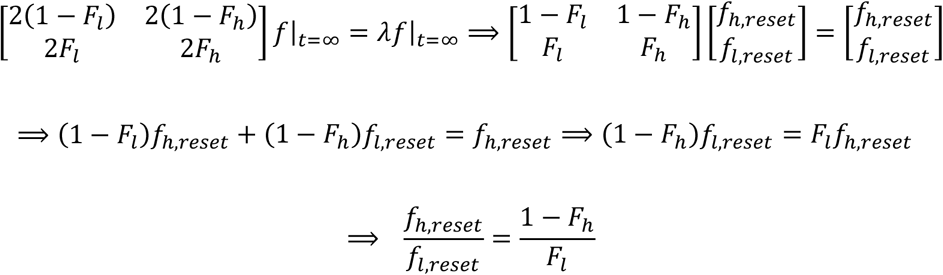

Then, analogously to the eigenvalue in the case of the standard MEN-model, we write the growth of the total population per generation in the case of feedback suppression:

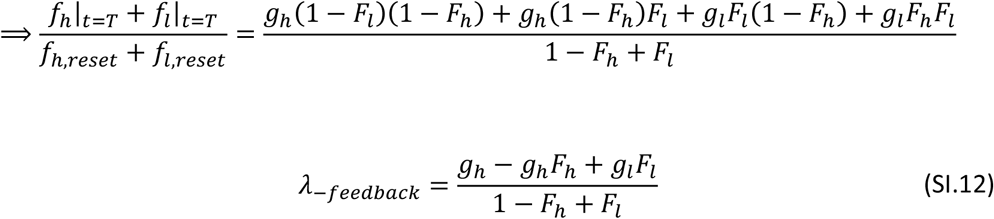

We compare this to the growth (eigenvalue) in the standard MEN-model case (including feedback):

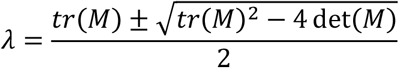

We can show that this *λ* is always at least as large as (*g_h_* – *g_h_F_h_* + *g_l_F_l_*)/(1 – *F_h_* + *F_l_*). In that case, the following identity must hold:

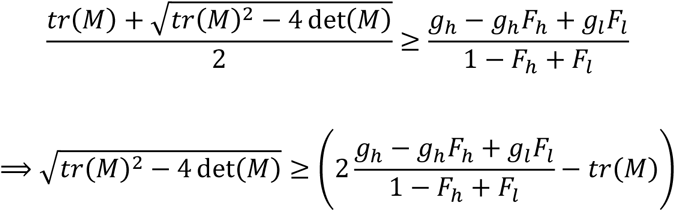

The left-hand side must be larger than zero, as *λ* must be real. If 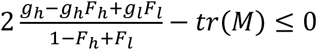, then the identity holds. When 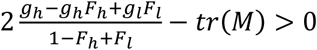, it is not immediately clear the identity holds. We need to check this case, where we can square both sides of the identity without flipping the ≥ sign:

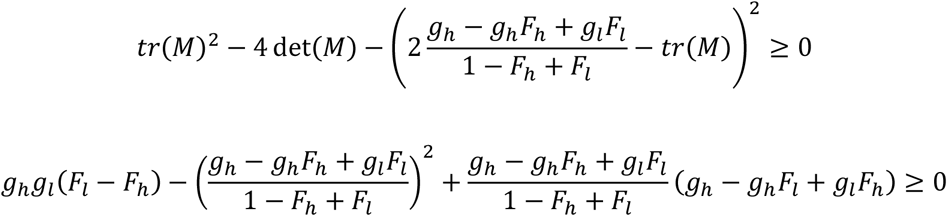

Placing terms under the same denominator:

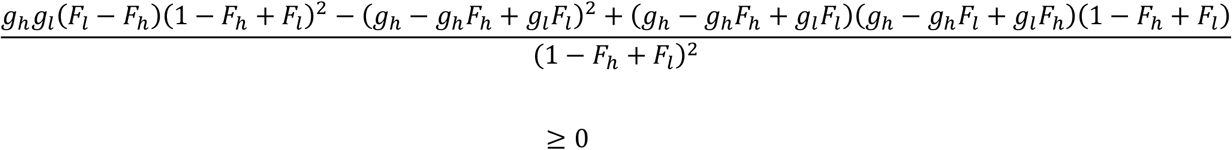

When analyzing first the numerator alone, we see many terms cancel out:

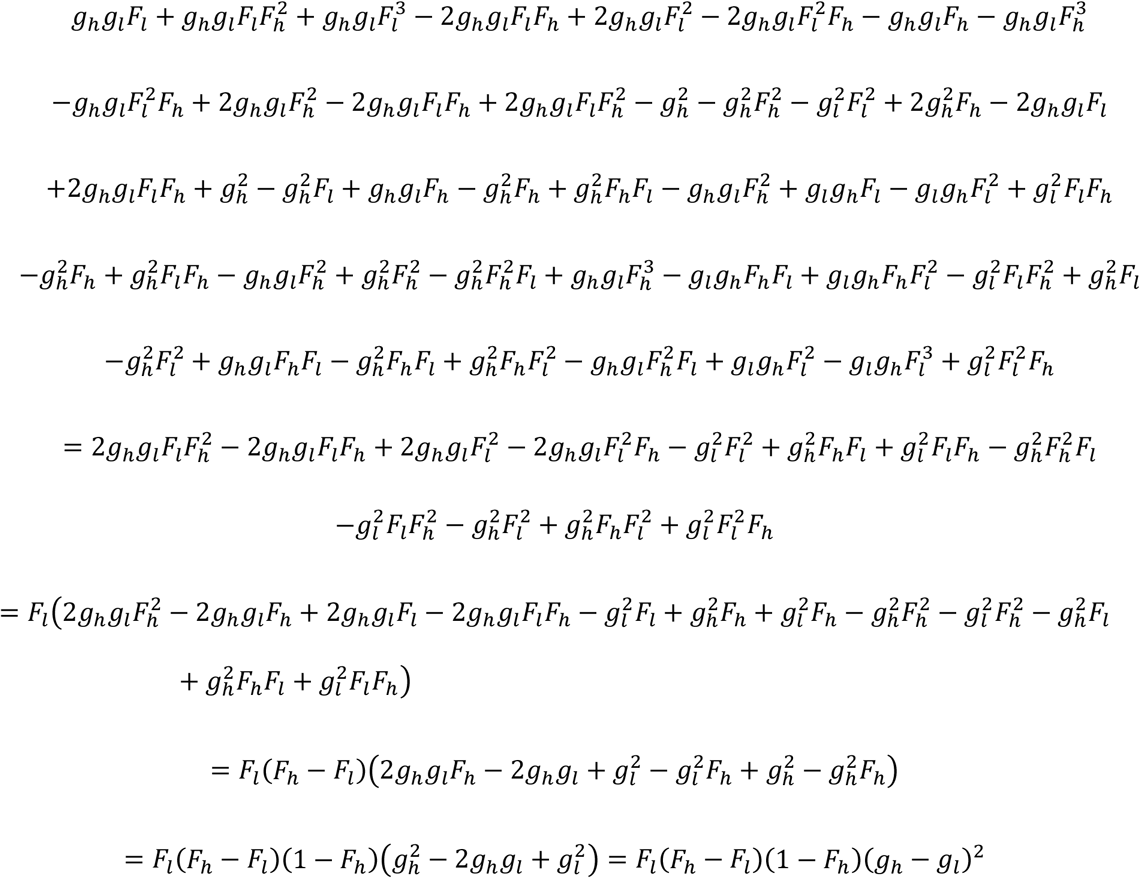

The identity to prove thus reduces to:

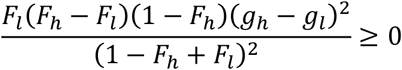

We can see this will always hold, as *F_h_* > *F_l_* > 0. The equality is only obtained when the progeny landscape is flat (*g_l_* = *g_h_*) or when there is no noise (*F_l_* = *F_h_*), or at least effectively no noise to use for switching of states (*F_l_* = 0 or 1 – *F_h_* = 0).

### Theoretical fitness landscapes

Figure 2 demonstrated the fitness landscapes for the two extreme cases of an essential and toxic gene. However, many genes fall in between these two cases (Figure 3-figure supplement 1B). Therefore, Figure 2-figure supplement 1 shows the fitness landscapes for mildly beneficial and mildly toxic genes, and with and without transgenerational feedback. In any case, we see how noise smoothens the landscape.

Additionally, to illustrate how the landscape is smoothened for realistic landscapes, we consider the landscape sharpness that we typically encounter from MEN-model fits on empirical landscapes of (Keren et al., 2016) (see Appendix 1-figure 1C). We define a width to represent the smoothing as the expression range spanning 10% to 90% of the fitness transition between the worst and best state (see in Figure 2-figure supplement 2A). This width will differ with and without feedback, and the ratio of widths between these two scenarios is plotted in Figure 2-figure supplement 2B for the possible landscape profile range from essential to toxic.

### Evaluation MEN-model fits on empirical fitness landscapes

The literature fitness landscapes of (Keren et al., 2016) as measured through an array of artificial promoters equate to the values of *λ_max_*/2 as a function of mean expression *μ*. To avoid the problems with the negative values for fluorescence relating to WT expression in (Keren et al., 2016), we combine the empirical landscapes with WT protein numbers of (Kulak et al., 2014). Furthermore, we also add essentiality data from (Cherry et al., 2012). The MEN-model fits improve the original fits of (Keren et al., 2016) (metric R^2^) for 63% of the cases. Because our parsimonious approach only requires 4 parameters (3 less than the original), adjusting our metric (Wherry, 1931) for this increase this percentages to 84% (see Fig. 2A). A similar percentage (85%) results from using the AIC (Akaike, 1974; Burnham & Anderson, 2004) as a metric.

The success of fitting fitness functions based on simple, sigmoidal progeny functions also seems in line with another study on a subset of these landscapes done in (Schmiedel, Carey, & Lehner, 2019), where a noise decomposition was also performed. There, authors demonstrate two recurrent noise-mean expression relation underlie most fitness landscapes. The essential/(mildly) beneificlal and (mildly) toxic landscape types we describe in Figure 2 and Figure 2-figure supplement 1 are interpretable as the principal topologies authors describe. Together with the fit metrics, this inspires trust in our approach, and we proceed to generate our model prediction of a realistic epistatic pattern.

Because the decomposition of observed fitness landscapes allows the decomposition into the progeny function and the noise component, we can also pose a different perspective to the observation that sharp fitness landscapes have lower noise levels (Keren et al., 2016). Remarkably, combining the fitted progeny sharpness *k* with noise levels that natively correspond to the respective genes (Chong et al., 2015) does not show a significant correlation between the two (Spearman *ρ* = 0.08 (p-value 0.59), N=40, see also Appendix 1-figure 1C). This test only includes genes with a known noise level in (Chong et al., 2015). No change in conclusion (Spearman *ρ* = −0.003 (p-value 0.99), N=24) follows by ignoring those *k* with large uncertainties (67% confidence interval > 10), which are mainly caused by relatively flat progeny landscapes. This prompts the hypothesis that observing a sharp fitness landscape implies low noise, rather than selection necessarily sharpening the fitness landscape due to low noise.

Moreover, we note that when we aggregate the noise of synthetic promoters used in (Keren et al., 2016) into a single noise level parameter, fits on the associated landscapes indicate relative high noise (Appendix 1-figure 1B). Concretely, for these genes the median value is 0.68, almost three times as large compared to 0.24 of (Chong et al., 2015). While we stress this fit parameter indicate an effective noise with diverse contributions, this indicates care must be taken with direct interpretations of the landscapes of (Keren et al., 2016). In particular, fitness costs of mutations that change noise level will otherwise be overestimated, as signaled by (Schmiedel et al., 2019), possibly contributing to the discrepancy discussed in the previous paragraph.

### Construction of DFEs

The parameter pool of the MEN-model fits is transformed to a new pool to generate a synthetic landscape pool (see Figure 3-figure supplement 1A). We removed the bias for essential genes (see Figure 3-figure supplement 1B) and combine this with a distribution of mutational effects (Figure 3-figure supplement 1C) as described in Simulation of representative DFEs. An example of such a DFE as function of background fitness is found in Figure 3-figure supplement 1C.

### Simulated DFE comparison to documented diminishing returns

To validate our simulated DFEs, we make use of the indel mutation type as our control. While removal of noise is inconsequential for this type, literature is available for an experimental DFE in yeast (Johnson et al., 2019). This transposon insertion induced DFE should theoretically be fairly comparable to our simulated indel DFE consisting of deletions of gene products. Yet, comparison with our simulations requires some filtering on the mutations. Many mutations will be almost neutral to within experimental resolution, In (Johnson et al., 2019), 64% of the mutations were deemed neutral, which roughly means that measured relative fitness effects of at most 0.4% are considered as neutral, which defines our neutrality threshold.

We then compare the slope of the observed diminishing returns pattern, a negative slope in mean mutational return as a function of background fitness. Because of the experimental design of (Johnson et al., 2019), we also exclude lethal mutations from our DFE, but only for the analyses on DFE statistics as a function of background fitness. Our slope of the mean fitness effect as function of background fitness is −0.29 (see Appendix 1-figure 2A), in reasonable accordance with figure 2E in (Johnson et al., 2019) where experimental slope for the mean is −0.18. By contrast, the slope in standard deviation of the DFE is not so well fitted (see Appendix 1-figure 2B), and only the sign of the trend is correct.

### Supplemental DFE results

We had seen in Figure 3 that the feedback does not necessarily imply a larger non-neutral mutation pool, even though this would have been possible theoretically (Figure 2). However, the landscape smoothing is clearly more noticeable when comparing essential against non-essential genes, see Figure 3-figure supplement 2. There, we see for the different DFEs the effect of essential genes on the non-neutral mutation pool. Again, the effect of feedback on the smoothing is not pronounced.

Figure 5 had focused on the point mutations. Duplications are not lethal, only in a rare case without feedback. Therefore, an analogous Figure 4 for duplication is not relevant, but an analogous Figure 5 is possible and shown in Figure 5-figure supplement 1.

Incidentally, we note that empirically a negative correlation exists between (log) mean expression and (log) noise level for certain expression levels (Bar-Even et al., 2006; Keren et al., 2015). Appendix 1-figure 3 shows the equivalent of Figure 3, Figure 4 and Figure 5, taking into account this negative correlation. However, we notice that this correlation has a negligible influence on our conclusions. Therefore, we consider for simplicity mutations that only change expression in the main text.

## Bibliography

Akaike, H. (1974). A new look at the statistical model identification. IEEE Transactions on Automatic Control, 19(6), 716–723.

Bank, C., Matuszewski, S., Hietpas, R. T., & Jensen, J. D. (2016). On the (un) predictability of a large intragenic fitness landscape. Proceedings of the National Academy of Sciences, 113(49), 14085–14090.

Bar-Even, A., Paulsson, J., Maheshri, N., Carmi, M., O’Shea, E., Pilpel, Y., & Barkai, N. (2006). Noise in protein expression scales with natural protein abundance. Nature Genetics, 38(6), 636–643.

Brauns, F., de la Cruz, L. M. I., Daalman, W. K.-G., de Bruin, I., Halatek, J., Laan, L., & Frey, E. (2020). Adaptability and evolution of the cell polarization machinery in budding yeast. bioRxiv. https://doi.org/10.1101/2020.09.09.290510

Burnham, K. P., & Anderson, D. R. (2004). Multimodel inference: understanding AIC and BIC in model selection. Sociological Methods & Research, 33(2), 261–304.

Cai, L., Friedman, N., & Xie, X. S. (2006). Stochastic protein expression in individual cells at the single molecule level. Nature, 440(7082), 358–362. https://doi.org/10.1038/nature04599

Cerulus, B., New, A. M., Pougach, K., & Verstrepen, K. J. (2016). Noise and Epigenetic Inheritance of Single-Cell Division Times Influence Population Fitness. Current Biology, 26(9), 1138–1147. https://doi.org/10.1016/j.cub.2016.03.010

Chechik, G., Oh, E., Rando, O., Weissman, J., Regev, A., & Koller, D. (2008). Activity motifs reveal principles of timing in transcriptional control of the yeast metabolic network. Nature Biotechnology, 26(11), 1251–1259.

Cherry, J. M., Hong, E. L., Amundsen, C., Balakrishnan, R., Binkley, G., Chan, E. T., … Wong, E. D. (2012). Saccharomyces Genome Database: the genomics resource of budding yeast. Nucleic Acids Research, 40(D1), D700–D705. https://doi.org/10.1093/nar/gkr1029

Chong, Y. T., Koh, J. L. Y., Friesen, H., Kaluarachchi Duffy, S., Cox, M. J., Moses, A., … Andrews, B. J. (2015). Yeast Proteome Dynamics from Single Cell Imaging and Automated Analysis. Cell, 161(6), 1413–1424. https://doi.org/10.1016/j.cell.2015.04.051

Chou, H.-H., Chiu, H.-C., Delaney, N. F., Segrè, D., & Marx, C. J. (2011). Diminishing returns epistasis among beneficial mutations decelerates adaptation. Science, 332(6034), 1190–1192.

Christiano, R., Nagaraj, N., Fröhlich, F., & Walther, T. C. (2014). Global Proteome Turnover Analyses of the Yeasts S. cerevisiae and S. pombe. Cell Reports, 9(5), 1959–1965. https://doi.org/10.1016/j.celrep.2014.10.065

Coomer, M. A., Ham, L., & Stumpf, M. P. H. (2022). Noise distorts the epigenetic landscape and shapes cell-fate decisions. Cell Systems, 13(1), 83–102.e6. https://doi.org/10.1016/j.cels.2021.09.002

Curk, T. (2016). Modelling multivalent interactons (PhD Thesis). University of Cambridge.

Daalman, W. K.-G., Sweep, E., & Laan, L. (2021). A tractable bottom-up model of the yeast polarity genotype-phenotype map for evolutionary relevant predictions. bioRxiv, 2020.11.09.374363. https://doi.org/10.1101/2020.11.09.374363

Dhondt, I., Petyuk, V. A., Bauer, S., Brewer, H. M., Smith, R. D., Depuydt, G., & Braeckman, B. P. (2017). Changes of Protein Turnover in Aging *Caenorhabditis elegans*. Molecular & Cellular Proteomics, 16(9), 1621–1633. https://doi.org/10.1074/mcp.RA117.000049

Diaz-Uriarte, R., & Vasallo, C. (2019). Every which way? On predicting tumor evolution using cancer progression models. PLoS Computational Biology, 15(8), e1007246.

Du, X., King, A. A., Woods, R. J., & Pascual, M. (2017). Evolution-informed forecasting of seasonal influenza A (H3N2). Science Translational Medicine, 9(413).

Duveau, F., Vande Zande, P., Metzger, B. P., Diaz, C. J., Walker, E. A., Tryban, S., … Wittkopp, P. J. (2021). Mutational sources of trans-regulatory variation affecting gene expression in Saccharomyces cerevisiae. eLife, 10. https://doi.org/10.7554/eLife.67806

Farslow, J. C., Lipinski, K. J., Packard, L. B., Edgley, M. L., Taylor, J., Flibotte, S., … Bergthorsson, U. (2015). Rapid Increase in frequency of gene copy-number variants during experimental evolution in Caenorhabditis elegans. BMC Genomics, 16(1). https://doi.org/10.1186/s12864-015-2253-2

Ferrell, J. E., & Ha, S. H. (2014a). Ultrasensitivity part I: Michaelian responses and zero-order ultrasensitivity. Trends in Biochemical Sciences, 39(10), 496–503. https://doi.org/10.1016/j.tibs.2014.08.003

Ferrell, J. E., & Ha, S. H. (2014b). Ultrasensitivity part III: cascades, bistable switches, and oscillators. Trends in Biochemical Sciences, 39(12), 612–618. https://doi.org/10.1016/j.tibs.2014.10.002

Ferrell, J. E., Jr, & Ha, S. H. (2014). Ultrasensitivity part II: multisite phosphorylation, stoichiometric inhibitors, and positive feedback. Trends in Biochemical Sciences, 39(11), 556–569. https://doi.org/10.1016/j.tibs.2014.09.003

Fish, D. A., Brinicombe, A. M., Pike, E. R., & Walker, J. G. (1995). Blind deconvolution by means of the Richardson–Lucy algorithm. JOSA A, 12(1), 58–65.

Fraser, H. B., Hirsh, A. E., Giaever, G., Kumm, J., & Eisen, M. B. (2004). Noise Minimization in Eukaryotic Gene Expression. PLoS Biology, 2(6), e137. https://doi.org/10.1371/journal.pbio.0020137

Fraser, H. B., Hirsh, A. E., Steinmetz, L. M., Scharfe, C., & Feldman, M. W. (2002). Evolutionary rate in the protein interaction network. Science, 296(5568), 750–752.

Friedman, N., Cai, L., & Xie, X. S. (2006). Linking stochastic dynamics to population distribution: an analytical framework of gene expression. Physical Review Letters, 97(16), 168302.

Giaever, G., Chu, A. M., Ni, L., Connelly, C., Riles, L., Véronneau, S., … Andre, B. (2002). Functional profiling of the Saccharomyces cerevisiae genome. Nature, 418(6896), 387–391.

Hodgins-Davis, A., Duveau, F., Walker, E. A., & Wittkopp, P. J. (2019a). Dataset for ‘Empirical measures of mutational effects define neutral models of regulatory evolution in Saccharomyces cerevisiae’. University of Michigan - Deep Blue. Retrieved from https://doi.org/10.7302/0dvr-k169

Hodgins-Davis, A., Duveau, F., Walker, E. A., & Wittkopp, P. J. (2019b). Empirical measures of mutational effects define neutral models of regulatory evolution in Saccharomyces cerevisiae. Proceedings of the National Academy of Sciences, 116(42), 21085–21093.

Houri-Zeevi, L., Korem Kohanim, Y., Antonova, O., & Rechavi, O. (2020). Three Rules Explain Transgenerational Small RNA Inheritance in C. elegans. Cell, 182(5), 1186–1197.e12. https://doi.org/10.1016/j.cell.2020.07.022

Jablonka, E., & Szathmáry, E. (1995). The evolution of information storage and heredity. Trends in Ecology & Evolution, 10(5), 206–211.

Johnson, M. S., Martsul, A., Kryazhimskiy, S., & Desai, M. M. (2019). Higher-fitness yeast genotypes are less robust to deleterious mutations. Science, 366(6464), 490–493.

Keren, L., Hausser, J., Lotan-Pompan, M., Vainberg Slutskin, I., Alisar, H., Kaminski, S., … Segal, E. (2016). Massively Parallel Interrogation of the Effects of Gene Expression Levels on Fitness. Cell, 166(5), 1282–1294.e18. https://doi.org/10.1016/j.cell.2016.07.024

Keren, L., van Dijk, D., Weingarten-Gabbay, S., Davidi, D., Jona, G., Weinberger, A., … Segal, E. (2015). Noise in gene expression is coupled to growth rate. Genome Research, 25(12), 1893–1902. https://doi.org/10.1101/gr.191635.115

Khan, A. I., Dinh, D. M., Schneider, D., Lenski, R. E., & Cooper, T. F. (2011). Negative epistasis between beneficial mutations in an evolving bacterial population. Science, 332(6034), 1193–1196.

Kimura, M. (1983). The neutral theory of molecular evolution. Cambridge University Press.

Kleijn, I. T., Krah, L. H. J., & Hermsen, R. (2018). Noise propagation in an integrated model of bacterial gene expression and growth. PLOS Computational Biology, 14(10), e1006386. https://doi.org/10.1371/journal.pcbi.1006386

Kosuri, S., Goodman, D. B., Cambray, G., Mutalik, V. K., Gao, Y., Arkin, A. P., … Church, G. M. (2013). Composability of regulatory sequences controlling transcription and translation in *Escherichia coli*. Proceedings of the National Academy of Sciences, 110(34), 14024–14029. https://doi.org/10.1073/pnas.1301301110

Kryazhimskiy, S., Rice, D. P., Jerison, E. R., & Desai, M. M. (2014). Global epistasis makes adaptation predictable despite sequence-level stochasticity. Science, 344(6191), 1519–1522.

Kulak, N. A., Pichler, G., Paron, I., Nagaraj, N., & Mann, M. (2014). Minimal, encapsulated proteomic-sample processing applied to copy-number estimation in eukaryotic cells. Nature Methods, 11(3), 319–324. https://doi.org/10.1038/nmeth.2834

Kvitek, D. J., & Sherlock, G. (2011). Reciprocal Sign Epistasis between Frequently Experimentally Evolved Adaptive Mutations Causes a Rugged Fitness Landscape. PLoS Genetics, 7(4), e1002056. https://doi.org/10.1371/journal.pgen.1002056

Larrimore, K. E., & Rancati, G. (2019). The conditional nature of gene essentiality. Current Opinion in Genetics & Development, 58, 55–61.

Li, X., Lalić, J., Baeza-Centurion, P., Dhar, R., & Lehner, B. (2019). Changes in gene expression predictably shift and switch genetic interactions. Nature Communications, 10(1). https://doi.org/10.1038/s41467-019-11735-3

Loison, L. (2019). Canalization and genetic assimilation: Reassessing the radicality of the Waddingtonian concept of inheritance of acquired characters. Seminars in Cell & Developmental Biology, 88, 4–13. https://doi.org/10.1016/j.semcdb.2018.05.009

Mineta, K., Matsumoto, T., Osada, N., & Araki, H. (2015). Population genetics of non-genetic traits: Evolutionary roles of stochasticity in gene expression. Gene, 562(1), 16–21. https://doi.org/10.1016/j.gene.2015.03.011

Miton, C. M., & Tokuriki, N. (2016). How mutational epistasis impairs predictability in protein evolution and design. Protein Science, 25(7), 1260–1272.

Murray, A. W. (2020). Can gene-inactivating mutations lead to evolutionary novelty? Current Biology, 30(10), R465–R471.

Muschiol, D., Schroeder, F., & Traunspurger, W. (2009). Life cycle and population growth rate of Caenorhabditis elegans studied by a new method. BMC Ecology, 9(1), 14. https://doi.org/10.1186/1472-6785-9-14

Mutalik, V. K., Guimaraes, J. C., Cambray, G., Lam, C., Christoffersen, M. J., Mai, Q.-A., … Endy, D. (2013). Precise and reliable gene expression via standard transcription and translation initiation elements. Nature Methods, 10(4), 354–360. https://doi.org/10.1038/nmeth.2404

Nagar, N., Ecker, N., Loewenthal, G., Avram, O., Ben-Meir, D., Biran, D., … Pupko, T. (2021). Harnessing machine learning to unravel protein degradation in Escherichia coli. Msystems, 6(1), e01296–20.

Neher, R. A., & Bedford, T. (2015). nextflu: real-time tracking of seasonal influenza virus evolution in humans. Bioinformatics, 31(21), 3546–3548. https://doi.org/10.1093/bioinformatics/btv381

Panchy, N., Lehti-Shiu, M., & Shiu, S.-H. (2016). Evolution of Gene Duplication in Plants. Plant Physiology, 171(4), 2294–2316. https://doi.org/10.1104/pp.16.00523

Philippi, T., & Seger, J. (1989). Hedging one’s evolutionary bets, revisited. Trends in Ecology & Evolution, 4(2), 41–44.

Prinz, H. (2010). Hill coefficients, dose–response curves and allosteric mechanisms. Journal of Chemical Biology, 3(1), 37–44. https://doi.org/10.1007/s12154-009-0029-3

Rutherford, S. L., & Lindquist, S. (1998). Hsp90 as a capacitor for morphological evolution. Nature, 396(6709), 336–342.

Sailer, Z. R., & Harms, M. J. (2017). High-order epistasis shapes evolutionary trajectories. PLOS Computational Biology, 13(5), e1005541. https://doi.org/10.1371/journal.pcbi.1005541

Sánchez, Á., Vila, J. C., Chang, C.-Y., Diaz-Colunga, J., Estrela, S., & Rebolleda-Gomez, M. (2021). Directed evolution of microbial communities. Annual Review of Biophysics, 50, 323.

Sanjuán, R., & Elena, S. F. (2006). Epistasis correlates to genomic complexity. Proceedings of the National Academy of Sciences, 103(39), 14402–14405.

Schmiedel, J. M., Carey, L. B., & Lehner, B. (2019). Empirical mean-noise fitness landscapes reveal the fitness impact of gene expression noise. Nature Communications, 10(1). https://doi.org/10.1038/s41467-019-11116-w

Schoustra, S., Hwang, S., Krug, J., & de Visser, J. A. G. M. (2016). Diminishing-returns epistasis among random beneficial mutations in a multicellular fungus. Proceedings of the Royal Society B: Biological Sciences, 283(1837), 20161376. https://doi.org/10.1098/rspb.2016.1376

Shen, X., Song, S., Li, C., & Zhang, J. (2022). Synonymous mutations in representative yeast genes are mostly strongly non-neutral. Nature. https://doi.org/10.1038/s41586-022-04823-w

Simpson, G. G. (1953). The Baldwin effect. Evolution, 7(2), 110–117.

Smith, N. J., Walt, S. van der, & Firing, E. (2015). Magma, inferno, plasma and viridis colormaps. Retrieved from https://github.com/BIDS/colormap/blob/master/colormaps.py

Stark, C., Breitkreutz, B.-J., Reguly, T., Boucher, L., Breitkreutz, A., & Tyers, M. (2006). BioGRID: a general repository for interaction datasets. Nucleic Acids Research, 34(90001), D535–D539. https://doi.org/10.1093/nar/gkj109

Waddington, C. H. (1942). Canalization of development and the inheritance of acquired characters. Nature, 150(3811), 563.

Waddington, C. H. (1953). Genetic assimilation of an acquired character. Evolution, 118–126.

Wagner, G. P., Pavlicev, M., & Cheverud, J. M. (2007). The road to modularity. Nature Reviews Genetics, 8(12), 921–931. https://doi.org/10.1038/nrg2267

Wang, Z., & Zhang, J. (2011). Impact of gene expression noise on organismal fitness and the efficacy of natural selection. Proceedings of the National Academy of Sciences, 108(16), E67–E76. https://doi.org/10.1073/pnas.1100059108

Wapinski, I., Pfeffer, A., Friedman, N., & Regev, A. (2007). Natural history and evolutionary principles of gene duplication in fungi. Nature, 449(7158), 54–61. https://doi.org/10.1038/nature06107

Wherry, R. J. (1931). A new formula for predicting the shrinkage of the coefficient of multiple correlation. The Annals of Mathematical Statistics, 2(4), 440–457.

Xue, B., & Leibler, S. (2016). Evolutionary learning of adaptation to varying environments through a transgenerational feedback. Proceedings of the National Academy of Sciences, 113(40), 11266–11271. https://doi.org/10.1073/pnas.1608756113

Yu, H., Greenbaum, D., Lu, H. X., Zhu, X., & Gerstein, M. (2004). Genomic analysis of essentiality within protein networks. TRENDS in Genetics, 20(6), 227–231.

Yu, J., Xiao, J., Ren, X., Lao, K., & Xie, X. S. (2006). Probing gene expression in live cells, one protein molecule at a time. Science, 311(5767), 1600–1603.

Zhang, J. (2003). Evolution by gene duplication: an update. Trends in Ecology & Evolution, 18(6), 292–298. https://doi.org/10.1016/S0169-5347(03)00033-8

Zhang, Z., Qian, W., & Zhang, J. (2009). Positive selection for elevated gene expression noise in yeast. Molecular Systems Biology, 5(1), 299. https://doi.org/10.1038/msb.2009.58

Zomorrodi, A. R., & Segrè, D. (2016). Synthetic Ecology of Microbes: Mathematical Models and Applications. Journal of Molecular Biology, 428(5), 837–861. https://doi.org/10.1016/j.jmb.2015.10.019

